# Late lactation represents the main window for sow-to-piglet transmission of persistent gut strains

**DOI:** 10.64898/2026.09.23.753686

**Authors:** Maëlle Pomiès, Gabryelle Agoutin, Lucas Auer, Adrien Castinel, Olivier Bouchez, Caroline S. Achard, Hélène Quesnel, Géraldine Pascal, Sylvie Combes

## Abstract

The gut microbiota plays a key role in piglet health, and maternal microbial transmission may represent a promising lever to shape early-life microbiota and prevent post-weaning digestive disorders. This study aimed to better characterize sow-to-piglet microbiota transmission and persistence using a long-read metabarcoding approach targeting the 16S-ITS-23S region. Fecal samples (n = 204) were collected from 17 families, a family being as sow and three of her piglets, at multiple stages: late gestation (G110), early (L6) and late lactation (L28) for sows; early lactation (L6), late lactation (L28), and 5 days post-weaning for piglets. To approximate strain-level resolution, a putative strain (PS) approach was developed by clustering ASVs (n = 6064) affiliated with the same species based on abundance covariance (r ≥ 0.9), resulting in 4857 PS. Piglet microbiota progressively diversified during lactation and converged toward that of sow. In sows, 27 ± 6% of PS were persistent from late gestation to late lactation. In piglets, only 4.2 ± 2.5% of PS persisted from d6 to 5 days post-weaning. Persistent PS in piglets were mainly affiliated with *Limosilactobacillus reuteri* and *Lactobacillus amylovorus* followed with *Holdemanella porci* and *H. biformis, Lentihominibacter hominis* and *Dorea formicigenerans*. Shared PS were significantly higher within families than between unrelated pairs (p < 0.05). Maternal transmission peaked at the end of lactation (35 ± 7% at L28). Persistent transmitted PS represented 2.7 ± 1.6% (d6-post-weaning) and 15.4 ± 5.6% (d28-post-weaning). Early-transmitted persistent PS were mainly affiliated with *Limosilactobacillus reuteri, Lactobacillus amylovorus*, and *Paraeggerthella hominis*, whereas late-transmitted persistent PS were associated with *Prevotella* spp., *Sphaerochaeta globosa*, and *Bariatricus comes*. These findings highlight the significance of maternal transmission in shaping the post-weaning microbiota and identify late lactation as a critical window for microbiota transfer.

## Introduction

On most pig farms, piglets are weaned at around 3-5 weeks of age. Weaning is marked by profound changes, including a change in diet, separation from the sow, introduction to a new environment, and cohabitation with new individuals. These disturbances often result in a temporary reduction in feed intake, intestinal inflammation, and an imbalance in the intestinal microbiota, (dysbiosis). These alterations are major risk factors for enteric disorders, particularly post-weaning diarrhoea (Gresse et al., 2017) that are often treated with antibiotics, contributing to the development of antibiotic resistance. The close relationship between human, animal and environmental health is now widely recognised, emphasising the importance of limiting antibiotic therapy and developing strategies to promote the health of farm animals from an early age. In this context, attention has been paid to identifying breeding strategies during gestation and/or lactation in order to help piglets withstand the challenges of weaning (Blavi et al., 2021). Targeted nutritional strategies for pregnant and lactating sows can improve their metabolic and health status and increase the immunological and nutritional quality of colostrum and milk (Hurley, 2015; Quesnel & Farmer, 2019).

Over the past decade, the significance of the digestive microbiota as a key factor in the health and development of piglets before and after weaning has been increasingly recognised (Gresse et al., 2017). Indeed, the intestinal microbiota has been shown to play a central role in the health development of young mammals. Specifically, it is involved in the metabolism of certain foods and the production of essential compounds, in addition to its role in defending against pathogens. As with all mammals, the digestive tract of pigs is colonised by more than three million microorganisms representing hundreds of species (Sender et al., 2016). This colonisation really begins at birth. At this time, the newborn is exposed to a variety of microbes that form a metacommunity. This metacommunity consists of microbe communities found in the vaginal and digestive tracts of the mother, as well as on her skin and in the immediate environment of the newborn (Dominguez-Bello et al., 2010). A random sample of bacteria from this metacommunity with high dispersal capabilities and rapid growth generally colonises ecological niches first and further facilitate the success of later colonizers (Connell & Slatyer, 1977; Christian et al., 2015). It is evident that the characteristics of the microbiota community in newborns are influenced by the diversity, relative abundance, and spatial distribution of the colonising bacteria in the metacommunity (Curtis & Sloan, 2004).

Transmitting targeted maternal microbiota to piglets could be an effective way of modulating their microbiota, thereby optimising their health and development. Jiménez et al., (2008) administered a genetically marked strain of *Enterococcus fecium* to pregnant mice. The same strain was subsequently found in the meconium of their offspring obtained by caesarean section, demonstrating that bacteria from the maternal gastrointestinal tract can be transmitted to newborns during pregnancy. In pigs, direct vertical transmission was first evidenced by Buddington et al., (2010), who supplemented sows with probiotics (*Lactobacillus acidophilus* or *Bifidobacterium lactis*) during late gestation and detected the strains in their offspring. Combining metabarcoding analyses and source tracking algorithm, it has been estimated that approximately 58.5% of the infant microbiota in humans originates from maternal sources, including the vaginal and gastrointestinal tracts as well as the skin (Bogaert et al., 2023). Applying the same approach in pigs, Liu et al., (2019) reported that sow-derived fecal microbes account for 7-20% of the piglet’s large intestinal microbiota between days 7 and 35. However, these last two studies do not allow conclusions to be drawn about the species or even the strains that are preferentially transmitted. In human, metagenomic tracking of maternal-derived strains has shown that those originating from the maternal gut are more persistent and exhibit enhanced colonization in the infant intestine compared to other sources (Ferretti et al., 2018; Mitchell et al., 2020). However tracking strains using metagenomics sequencing is complexe and time consuming and do not cope with high-throughput analysis. While the exploration of microbiota has long relied on metabarcoding sequencing analysis of short fragments (500 bp), the latest advances in sequencing now provide access to the complete sequence of the 16S rRNA gene and even 16S-ITS-23S ribosomal gene operon (approximately 4,500 bp) (Karst et al., 2021). Recently, whole 16SrDNA gene sequencing was used to improved the characterisation of insect gut microbiota and track the establishment of the specific isolates (Mason et al., 2025).

The aim of the study was to evaluate the transmission of microbiota from sows to their offspring based on HiFi PacBio long sequencing reads from metabarcoding using the complete 16S-ITS-23S ribosomal gene operon as a target marker. To this end, we studied the fecal microbiota of 17 pig families (each consisting of one sow and three piglets) from the end of gestation to the early post-weaning period. The data presented herein were produced using the experimental study conducted by Le Floc’h et al., (2022). We tracked the persistence and transmission of covariant ASVs within a taxonomic species that we named putative strains.

## Material and Methods

### Animals and experimental design

The experiment was carried out at INRAE (UE3P, Saint-Gilles, France, doi: 10.15454/1.5573932732039927E12), in compliance with the Directive 2010/63/UE on animal experimentation. The experimental protocol was approved by the regional Ethics Committee in Animal Experiment of Rennes (France) and by the French Ministry of Higher Education, Research and Innovation (authorization APAFIS#11015-2017080716549316). This work was done with the same experimental design used to study the effect of live yeast supplementation in sow diet during gestation and lactation on sow and piglet fecal microbiota, health, and performance (Le Floc’h et al., 2022). Briefly, 48 Landrace x Large White sows and their litter were used in 4 batches. Sows were inseminated with semen from Piétrain boars. At 28 days of gestation (G28), sows were distributed into two dietary treatments: Control and SB. Sows in the SB group were fed the same standard gestation and then lactation diet as the Control sows but with the addition of *Saccharomyces cerevisiae var. boulardii* CNCM I-1079 (Levucell SB; Lallemand SAS, France) as live yeast cells (minimum concentration of 1 × 10^10^ colony-forming unit [CFU/g] added at 100 g/ton [1 × 10^9^ CFU/kg of feed]). From G28, sows were housed in groups of six in a pen with concrete floor (5 × 3.5 m) covered with wood hulls. At 106 days of gestation, sows were moved to the farrowing room and were kept in individual farrowing crates thereafter. During gestation, and until the day of farrowing, sows were fed a conventional gestation diet. From day 1 of lactation (day 0 of lactation being the day of farrowing), sows were fed a conventional lactation diet. The piglets were offered a conventional prestarter feed during the fourth week of lactation (for detailed feed composition see (Le Floc’h et al., 2022). Piglets were weaned at 28 days of lactation (L28) and vaccinated against porcine type 2 circovirus and Mycoplasma hyopneumoniae (Porcilis PCV M Hyo, MSD Santé Animale, France). No antibiotics were preventively provided through the diet or water. At weaning, piglets were transferred into a postweaning unit and group-housed in pens of 9 to 11 piglets. Each pen housed piglets from one experimental treatment only (Control or SB). Piglets with a similar range of body weight from at least four litters were mixed in the same pen (Table S1 https://doi.org/10.57745/3ZJXTI). Moreover, piglets were transferred into pens that were not cleaned after the departure of the previous batch and the ambient temperature was set at 24°C at the pig arrival in the postweaning building before being progressively increased until 28°C. They were offered the prestarter feed for the first 5 days (W5).

### Feces sampling, DNA extraction and long-read 16S-ITS-23S rRNA operon HiFi sequencing

Feces were collected from sows on day 110 of gestation (G110), 6, and 28 days of lactation (L6 and L28 respectively) and from three female piglets per litter on L6, L28, and 5 days after weaning (W5) after rectal stimulation (Figure 1). DNA was extracted from 40 to 60 mg of fecal material using the ZR-96 Soil Microbe DNA Kit (Zymo Research, Freiburg, Germany) according to the instructions of the manufacturer. A 15-minute bead-beating step at 30 Hz was performed using a Retsch MM400 mixer. The 16S-ITS-23S ribosomal operon was amplified by PCR using primers 16S-27F: AGRGTTYGATYMTGGCTCAG and 23S-2241R: ACCRCCCCAGTHAAACT. Amplicons were generated following the BOA (Barcoded Overhang Adapters) protocol. Sequencing was performed on a PacBio Sequel II system using HiFi long-read technology in the Genomic and Transcriptomic Platform (GeT-PLAGE, INRAE, Toulouse, France). The quality of the sequences and the read length distribution (raw FASTQ files) were assessed using FastQC v0.12.1, MultiQC v1.27.1 (Ewels et al., 2016), and seqkit v2.9.0 (Shen et al., 2016). Data selection was based on family completeness (availability of data for both sow and piglet at each sampling time point) and a minimum sequencing depth of 10000 reads per sample. Among the 48 pig families, 33 had complete sampling, and among the later 17 were retained based on sequence quality, resulting in a total of 204 fecal samples, 51 from sows and 153 from piglets Of the 17 retained families, 5 belonged to the Control group and 12 to the Yeast-supplemented group. As the primary objective of this analysis was to characterize mother-to-piglet bacterial transmission rather than treatment effects, treatment group was not included as a factor in the subsequent analyses. Animal metadata are provided in supplementary Table S1 (https://doi.org/10.57745/3ZJXTI) and sequences were deposited in European Nucleotide Archive (ENA): accession number is PRJEB113713.

**Figure 1:**
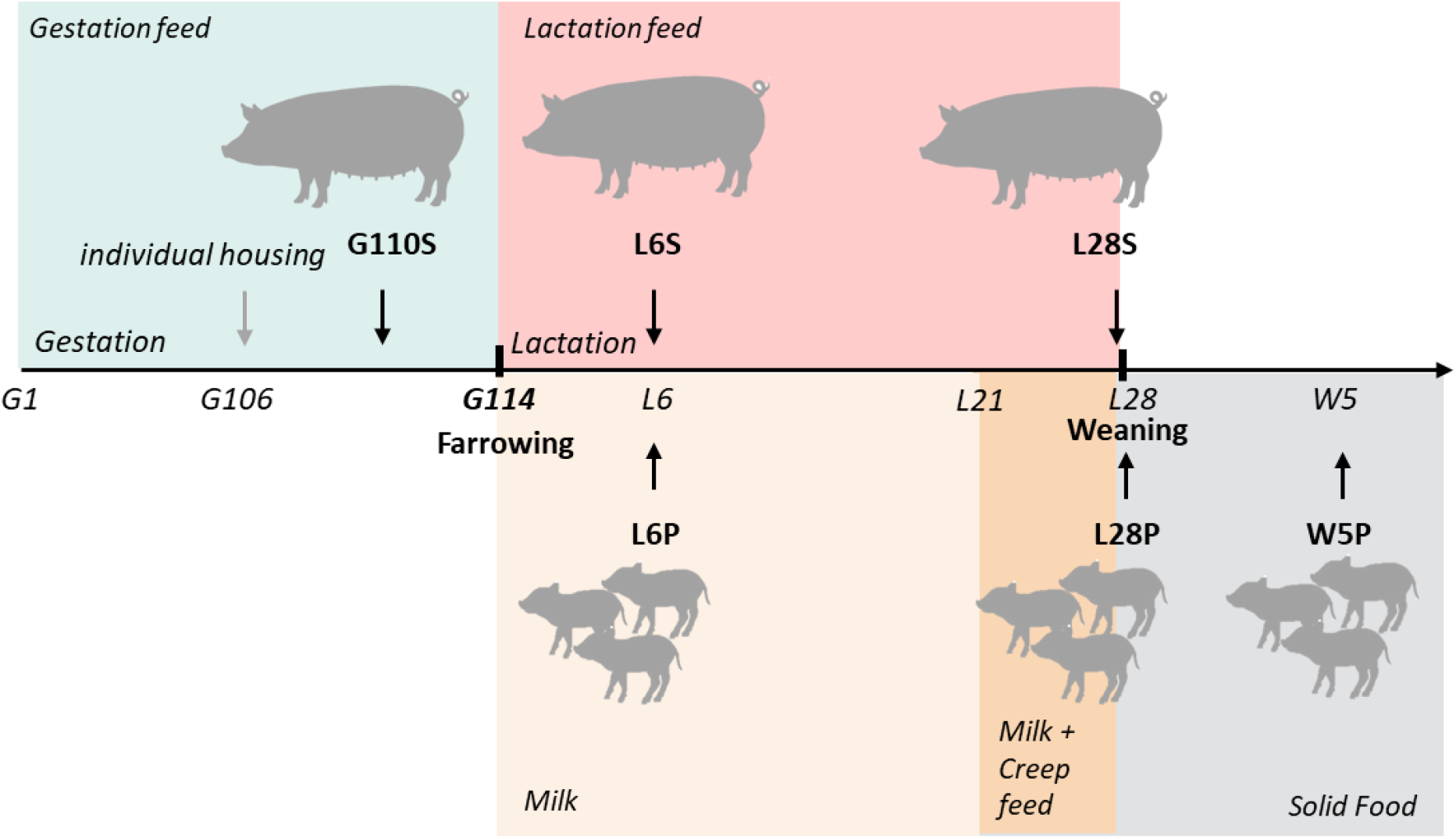
Experimental design. G stand fo gestation stage, L for lactation stage and W for weaning. G110S, L6S, and L28S correspond to sow fecal samples collected at the end of gestation (day 110), at the beginning of lactation (day 6), and at the end of lactation (day 28, weaning), respectively; L6P, L28P, and W5P correspond to piglet fecal samples collected at 6 and 28 days of suckling and 5 days after weaning, respectively. (n=17 sows, n= 51 piglets and 3 piglets per sow).

### Bioinformatics analysis of long-read 16S-ITS-23S rRNA operon amplicon

A total of 11,550,265 reads were analyzed using FROGS v.5.1 software (Escudié et al., 2018). We successively used the read_processing, remove_chimera, cluster_filter, taxonomic_affiliation, normalisation, taxonomic_affiliation and affiliation_filters tools with the parameters shown in Table 1. Command lines are available on https://forge.inrae.fr/geraldine.pascal/sow-to-piglet-data-article Before normalisation, the mean number of reads per sample in pig fecal sample was 46904 (min: 9913 - max: 224975) with an average length of 4200 ± 171 base paires. The 16S-ITS-23S ASV data were normalized to 10000 sequences per sample. Taxonomic affiliation was performed using 16S-ITS-23S-DB, a FROGS database (https://doi.org/10.57745/IRLQI1). ASVs with taxonomic assignments supported by less than 90% identity to a sequence in the 16S-ITS-23S-DB database were labeled with the attribute “weak-affiliation” to distinguish them from ASVs with more reliable taxonomic affiliations. Five fecal samples were extracted and sequenced in duplicate to visualized results repeatability (Supplemental Figure S1; https://forge.inrae.fr/geraldine.pascal/sow-to-piglet-data-article).

**Table 1.**
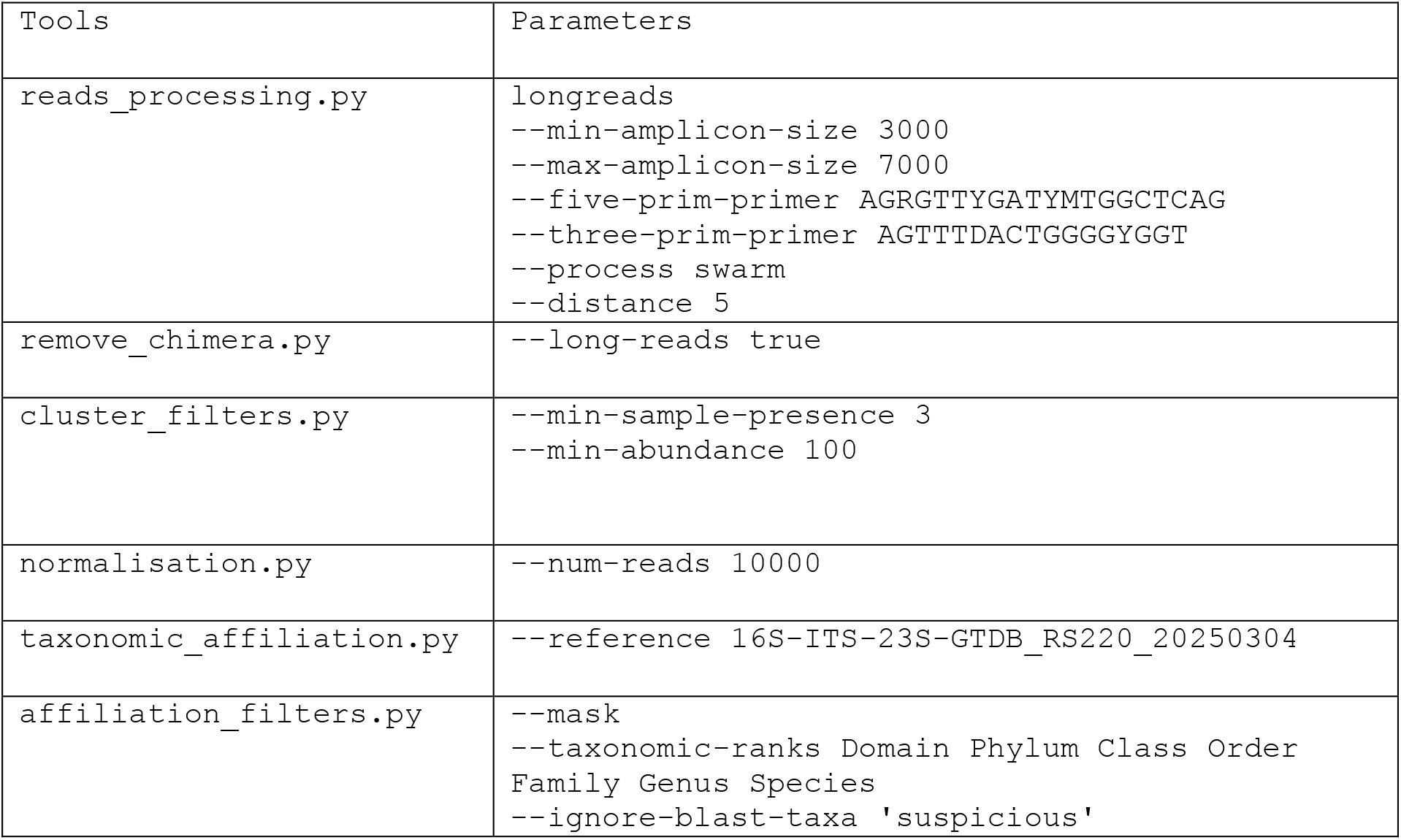
FROGS 5.1 parameters used for the bioinformatics analysis.

### Construction of putative strain from 16S-ITS-23S ASV clustering

The final step of the data processing workflow consisted of grouping 16S-ITS-23S ASVs belonging to the same species and showing covariation across all samples (ASVs with “weak-affiliation” and reliable affiliations treated separately). This step was performed using an R (version 4.2.1) script computing a Pearson correlation matrix based on the abundances of 16S-ITS-23S ASVs assigned to the same species. Hierarchical clustering was then applied to this correlation matrix using the “hclust” function, with correlation distance and the “average” linkage method. Groups were then defined using the cutree function, applying a correlation threshold of ≥ 0.9. This process yielded an abundance table of putative strains (PS) instead of individual 16S-ITS-23S ASVs.

### Statistical analysis

Statistical analyses were performed on 16S-ITS-23S derived putative strain (PS) using R software v4.2.1. The vegan package (Oksanen et al., 2026) was used to calculate the diversity indices and compute the Bray-Curtis dissimilarity matrix that was visualize using Principal Coordinates Analysis (PCoA). Pairwise differences between groups were assessed using PERMANOVA with the pairwiseAdonis package (Arbizu, 2017). Pairwise Wilcoxon tests were used to compare means of observed PS (richness) and Shannon diversity indices. Transmission and persistence of PS were assessed through co-occurrence analysis between relevant samples. Linear mixed models were used to compare transmission and persistence across developmental stages (gestation, lactation, weaning), with stage as a fixed effect, and piglet and sow as random effects to account for the hierarchical structure of the data. A second model was constructed to evaluate the effect of litter membership (same litter or not) on transmission. This model included two fixed effects, stage and litter membership (within or between), as well as their interaction. Post hoc comparisons were conducted, with Benjamini-Hochberg correction applied to control the false discovery rate.

**The number of persistent putative-strains (PS)** for individual *i* corresponds to the size of the intersection of all PS sets observed at each of the *n* time points *t* (G110, L6, L28 for sow and L6, L28 and W5 for piglets).

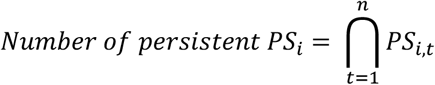

*i*: individual (e.g., sow or piglet)

*t*: time point (e.g., gestation, lactation, weaning)

*n*: total number of time points or stages

**Number of PS transmitted from sow i to her piglet j** corresponds to the intersection between all PS detected in sow *i* across *n* time points, and all PS detected in piglet *j* across m time points.

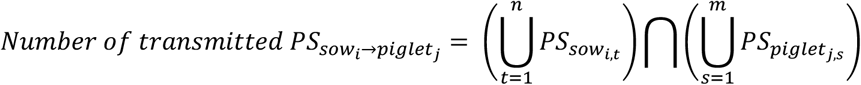

*i*: index of the sow

*j*index of one of her piglets

*t* = 1,…,n: time points or stages at which sow *i* was sampled

*s*= 1,…, m: time points or stages at which piglet *j* was sampled

**The number of PS transmitted by sow i to her piglet j, detected at least once during G110 or L6 in the sow and persisting in the piglet during all stages (L6, L28 and W5)** is given by the intersection between the union of PS from the sow during G110 and L6 and the intersection of PS from the piglet during L6, L28 and W5.

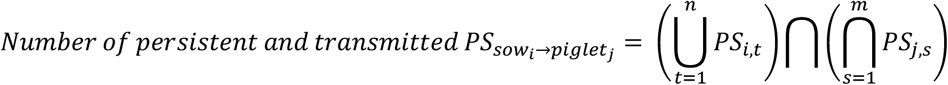

*i*: index of the sow

*j*index of one of her piglets

*t* = 1,…,n: dedicated time points at which sow i was sampled

*s* = 1,…, m: dedicated time points at which piglet j was sampled

Finally, persistence or transmission scores are calculated for each PS across all piglets at the different stages, as the sum of its absence (= 0) or presence (= 1), or as the sum of its absence of co-occurrence with the mother (= 0) or co-occurrence with the mother (= 1), respectively.

## Results

After sample rarefaction (n = 204 samples), sequences were clustered into 6064 ASVs and subsequently grouped into 4857 PS based on their species-level taxonomic assignment and covariance across the dataset. However, 67% of ASVs, accounting for 55% of the total sequences, could not be grouped, with 4,064 PS consisting of a single ASV (singletons).The proportion of PS that was assigned at the species level was 65.3%, which corresponds to 75.6% of the sequence count. This was based on > 90% sequence identity to reference sequences in the 16S-ITS-23S-DB database.

### Fecal bacterial community alpha and beta diversities evolve with time

As expected, fecal bacterial community diversity changed with age in piglets (Figures 2 and 3). Alpha-diversity increased over time, with a significant decrease in the Shannon index observed 5 days after weaning (Figure 2). In sows, alpha-diversity remained stable across stages (Figure 2), whereas beta-diversity only differed between late gestation and the two lactation stages (PEMANOVA R^2^ > 0.18, p < 0.05, Supplemental figure S2 https://forge.inrae.fr/geraldine.pascal/sow-to-piglet-data-article).

**Figure 2:**
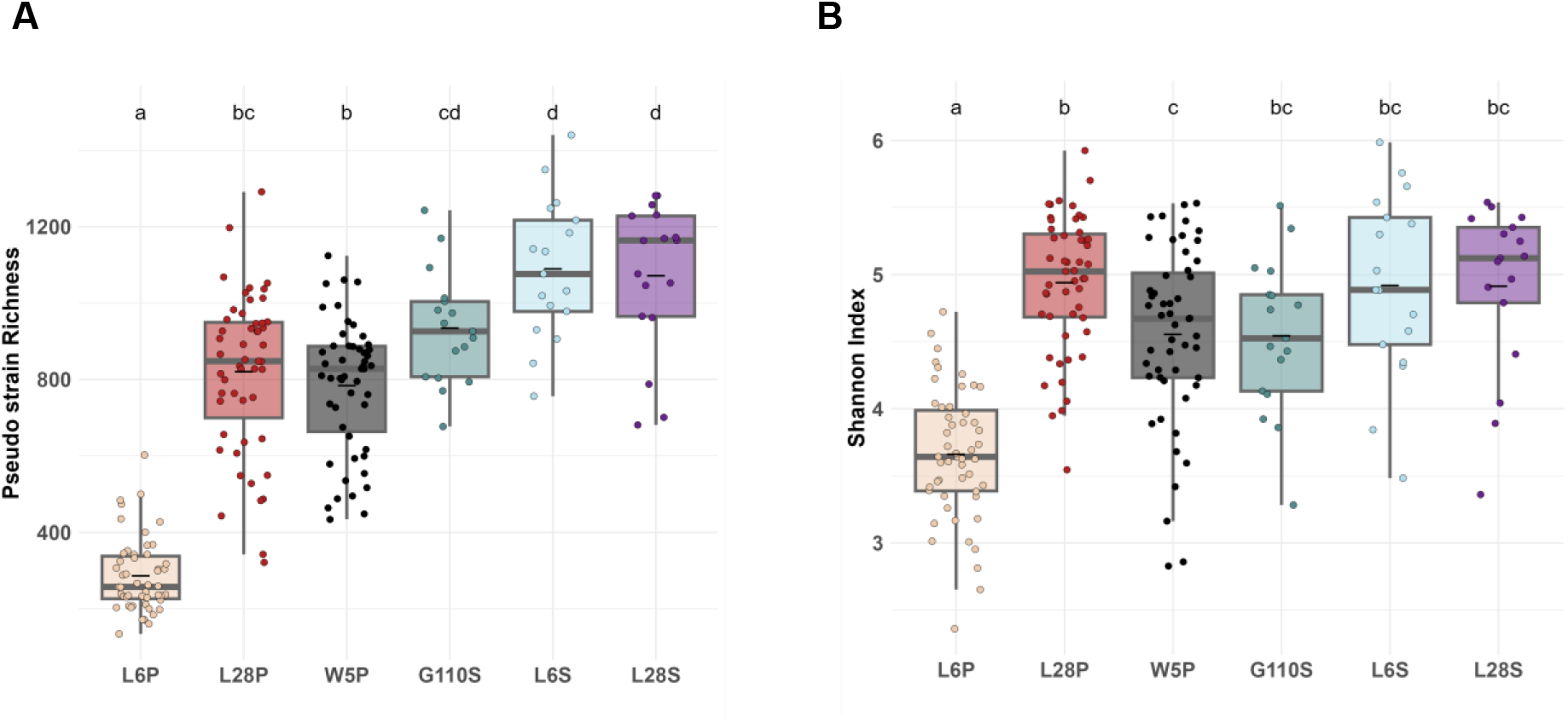
Fecal bacterial community α diversity in piglets and sows over time (n=17 sows, n= 51 piglets and 3 piglets per sow) on putative strain data. A. Putative strain richness, B. Shannon index. L6P, L28P, and W5P correspond to piglet fecal samples collected at 6 and 28 days of suckling and 5 days after weaning, respectively; G110S, L6S, and L28S correspond to sow fecal samples collected at the end of gestation (day 110), at the beginning of lactation (day 6), and at the end of lactation (day 28, weaning), respectively. Mean with different letter differ at p < 0.05.

**Figure 3:**
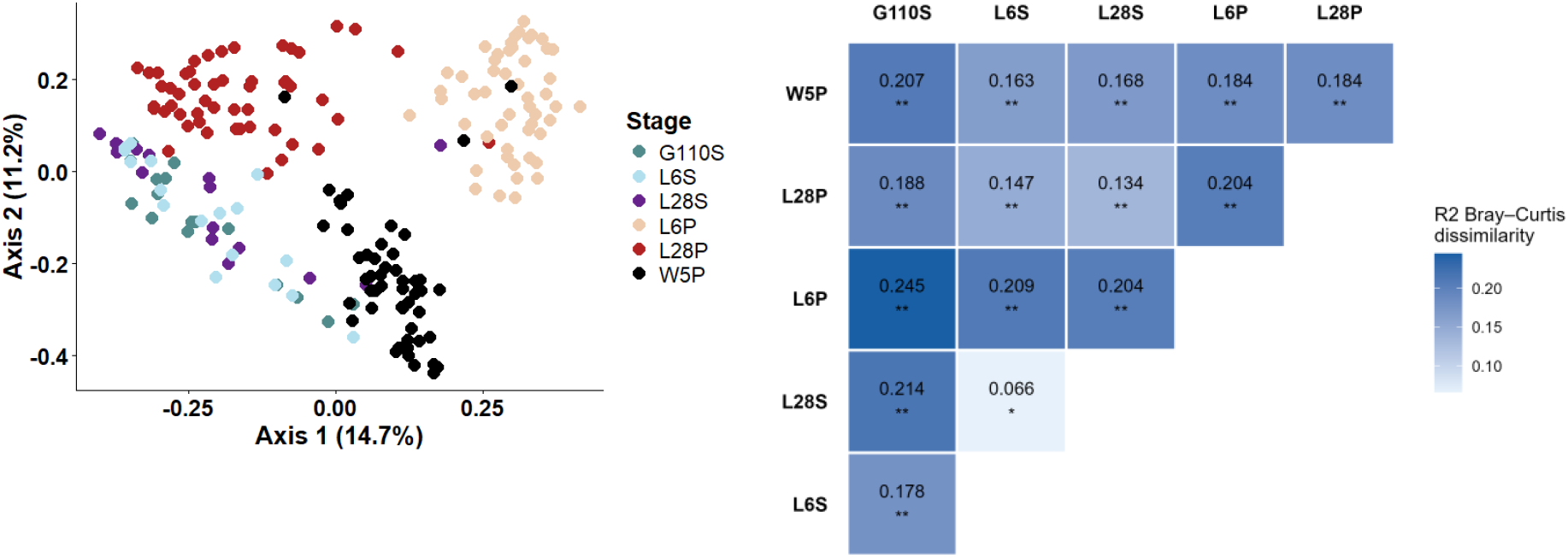
Fecal bacterial beta-diversity in piglets and sows over time on putative strain data (n=17 sows, n= 51 piglets and 3 piglets per sow) A. Principal coordinate analysis (PCoA) plot. Each point represents a single sow or piglet for a given time. B. Pairwise PEMANOVA R^2^ value (ADONIS). L6P, L28P, and W5P correspond to piglet fecal samples collected at 6 and 28 days of suckling and 5 days after weaning, respectively; G110S, L6S, and L28S correspond to sow fecal samples collected at the end of gestation (day 110), at the beginning of lactation (day 6), and at the end of lactation (day 28, weaning), respectively. * p < 0.05, ** p < 0.01

### Persistence of putative strain in sow and piglets

First, we assessed PS persistence by calculating the proportion of PS shared between samples at different stages within the same individual for sows and piglets independently (Figure 4). The proportion of persistent PS in sows (Figure 4A) between the end of gestation (G110) and the beginning of lactation (L6) did not differ significantly from that observed between the beginning (L6) and the end of lactation (L28) (p = 0.101). In contrast, the proportion of shared PS was significantly higher during the lactation period (L6-L28) than between the end of gestation and the end of lactation (G110-L28, p < 0.001). Consequently, the proportion of persistent PS from the end of gestation throughout lactation was the lowest, representing 27.2 ± 5.9% of the PS present at the end of lactation. Among the latest, a total of 15 PS were detected in all the 17 sows and 14 PS were affiliated at the species level with *Clostridium saudiense*, Intestinibacter bartlettii (2PS), *Lactobacillus amylovorus, Limosilactobacillus mucosae, Limosilactobacillus reuteri, Romboutsia D lituseburensis A, Romboutsia E sp900545985 (3PS), Romboutsia ilealis, Terrisporobacter mayombei, Terrisporobacter petrolearius, Turicibacter bilis*. Relative abundance patterns were PS-specific (Supplementary Figure S3 https://forge.inrae.fr/geraldine.pascal/sow-to-piglet-data-article). A decrease after parturition (G110) was observed for *Intestinibacter bartlettii* (significant for one PS), *Lactobacillus amylovorus, Limosilactobacillus mucosae*, and *Turicibacter bilis*. In contrast, an increase in relative abundance was observed for PS affiliated with *Limosilactobacillus reuteri, Terrisporobacter mayombei*, and *Terrisporobacter petrolearius*. Differential dynamics were also observed among PS within the same species, notably for *Romboutsia sp*., with distinct trends between PS. Finally, the highest relative abundances were observed at the end of lactation for eight PS, whereas three PS showed a decrease in relative abundance over the course of lactation (Supplementary Figure S3 https://forge.inrae.fr/geraldine.pascal/sow-to-piglet-data-article).

**Figure 4:**
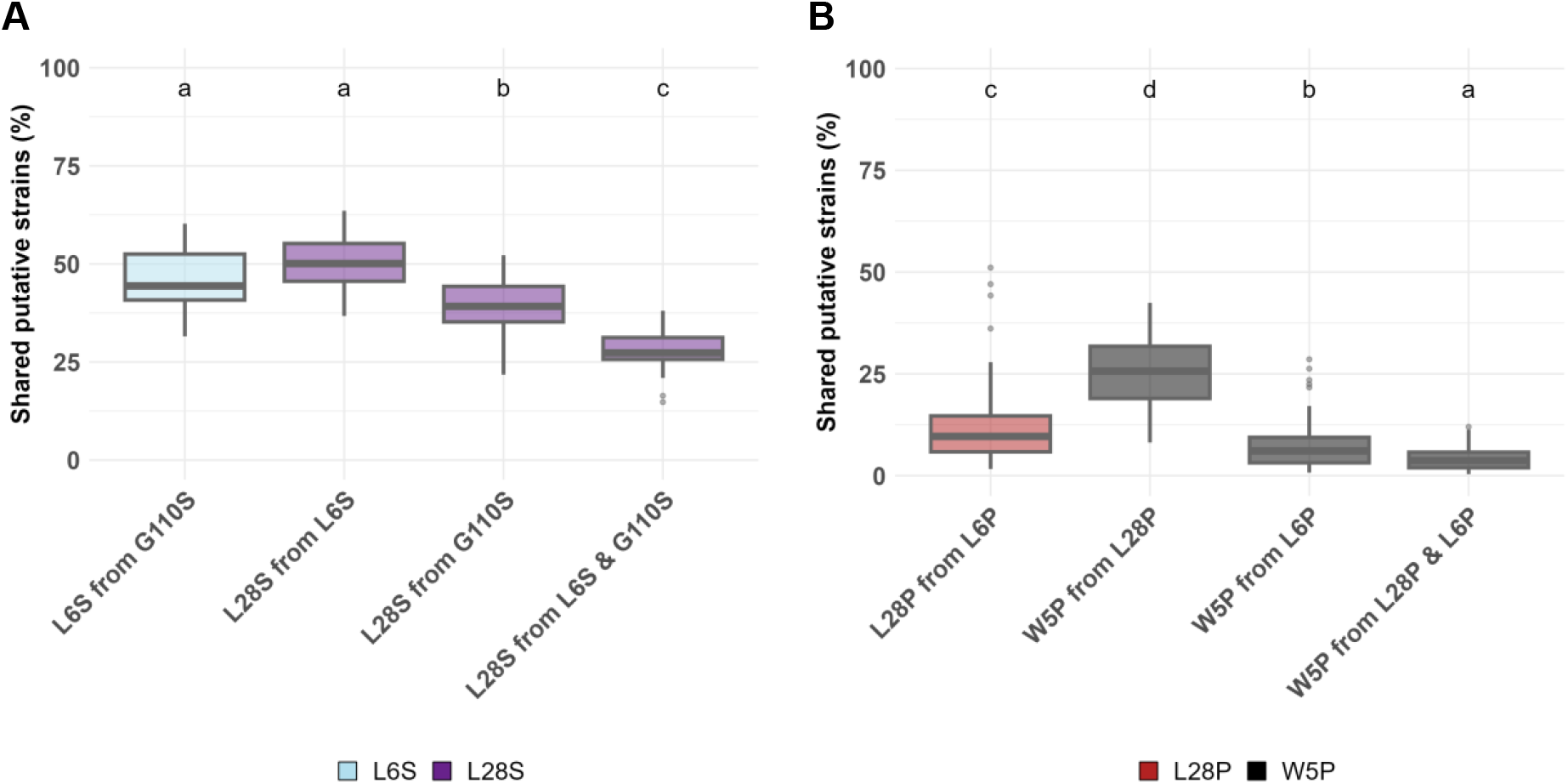
Persistence of putative strain between different sampling stages A. in sows and B in piglets within a family. G110S, L6S, and L28S correspond to sow fecal samples collected at the end of gestation (day 110), at the beginning of lactation (day 6), and at the end of lactation (day 28, weaning), respectively (n=17 sows). L6P, L28P, and W5P correspond to piglet fecal samples collected at 6 and 28 days of suckling and 5 days after weaning, respectively (n= 51 piglets with 3 piglets per sow). Different letters indicate a significant difference between means (p < 0.05).

In contrast, in piglets from the same litter, the proportion of persistent PS was lower than in sows, reaching 11.2 ± 7.8% throughout lactation and 25.1 ± 8.3% between the end of lactation and 5 days after weaning (Figure 4B). The highest proportion of persistent PS was observed between the end of lactation and weaning. Finally, only 4.1 ± 2.5 % of PS were detected both at the beginning and at the end of lactation and remained present until the 5 days after weaning. Among these, a total of 34 persistent PS were detected in all piglets of the 17 families (Figure 5A). PS with the highest persistent score across the 17 families (sum of absence = 0 or presence = 1) were affiliated at the species level with *Limosilactobacillus reuteri (3PS)* and *Lactobacillus amylovorus* followed with *Holdemanella porci* and *biformis, Lentihominibacter hominis* and *Dorea formicigenerans* (Figure 5B). PS persistence was strongly correlated with abundance (R = 0.74, p < 0.001, Supplementary Figure S4, https://forge.inrae.fr/geraldine.pascal/sow-to-piglet-data-article), although this relationship was no longer significant when restricted to the 34 PS detected in all piglets across the 17 families (R = 0.33, p = 0.053, Figure 5C)

**Figure 5:**
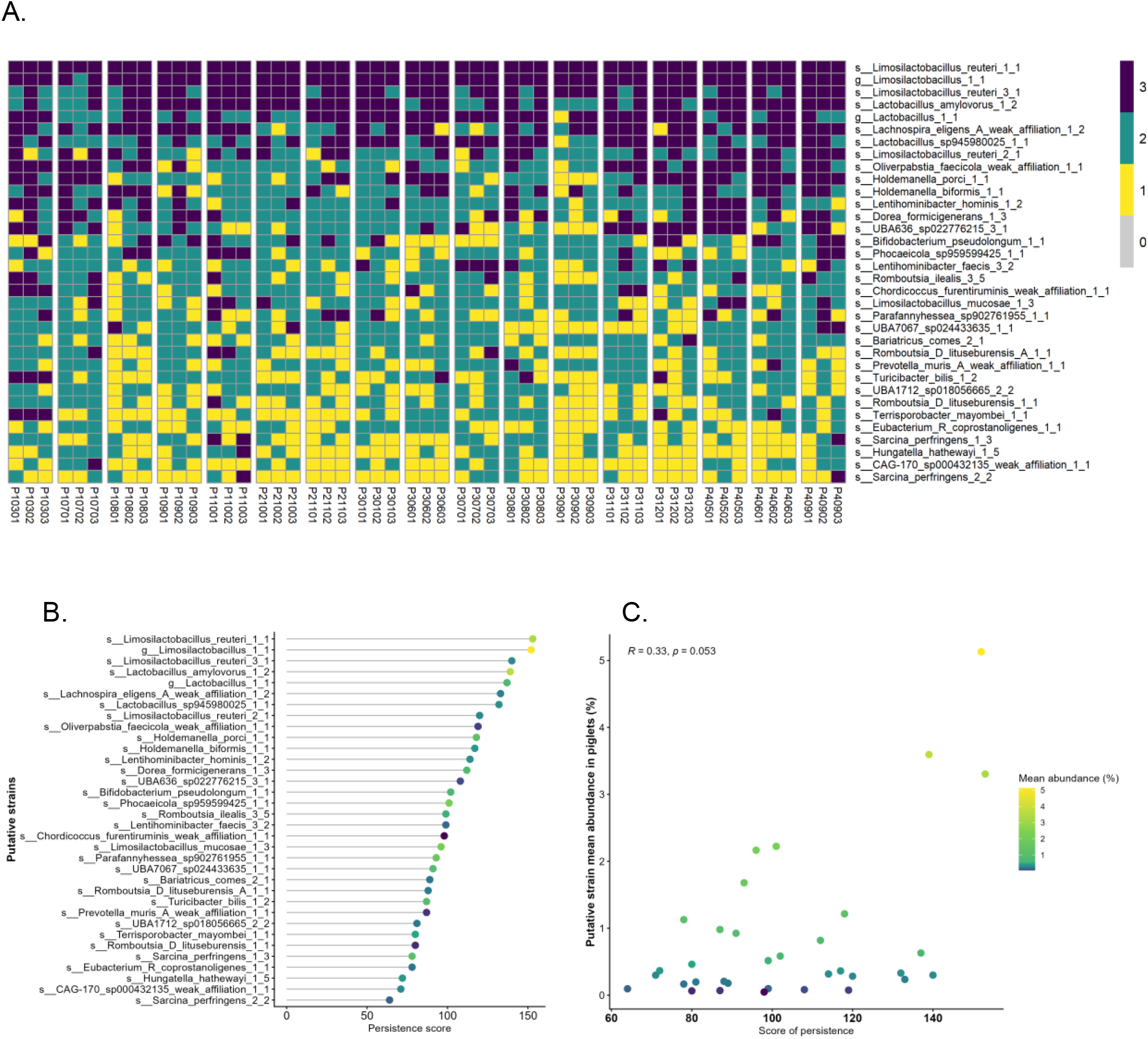
Persistent putative strains from lactation to five days after weaning in piglets. A. Heatmap showing piglet persitent putative strains. The rows correspond to the PS with taxonomic affiliations at the species (s_) or genus (g_) level. s_Limosilactobacillus_amylovorus_1_2 refers to PS 1 clustering two ASVs affiliated to *Limosilactobacillus amylovorus*. The columns represent the 51 piglets, grouped according by family (17 litters). Colors indicate persistence across stages: purple, persistent across all three stages; green, persistent across two stages; yellow, observed at a single stage. B. Lolipop plot showing the persistence score, defined as the sum of PS presence across all stages (n = 3) and all piglets (n = 51). C. Relationship between PS relative abundance and persistence score. Points are colored according to relative abundance (yellow: high; purple: low).

### Transmission of sow-derived gut putative strains in piglets

Then, we evaluated PS transmission from sow to her piglets by calculating the proportion of PS shared between samples at different stages within related sows and piglets. At the beginning of lactation (L6), the proportion of PS transmitted to the piglets from their sow (Figure 6A) did not differ significantly depending on whether the sow was at the end of gestation (G110S) or at the beginning of lactation (L6S). The transmitted PS proportion to piglets was significantly higher at the end of lactation (L28P) than at the beginning of lactation (L6P). Finally, piglets shared the most PS with their sow at the end of lactation (L28P from L28S). Five days after weaning (S5P), piglets also shared more PS with their sow at the end of lactation (L28S) than with the sow at the end of gestation (G110S) or at the beginning of lactation. However, the proportion of PS shared with the maternal microbiota at the end of lactation decreased after weaning, suggesting a reduced persistence of maternal-associated PS following the cessation of direct contact with the sow. We then determined whether PS were more frequently shared between piglets and their own sow than with unrelated sows (Figure 6B). The proportions of PS shared between piglets (P) and sows (S) were significantly higher within the same family than between unrelated piglets and sows, except when piglets were at the end of lactation (L28P) and sows were at the end of gestation (G110S).

**Figure 6:**
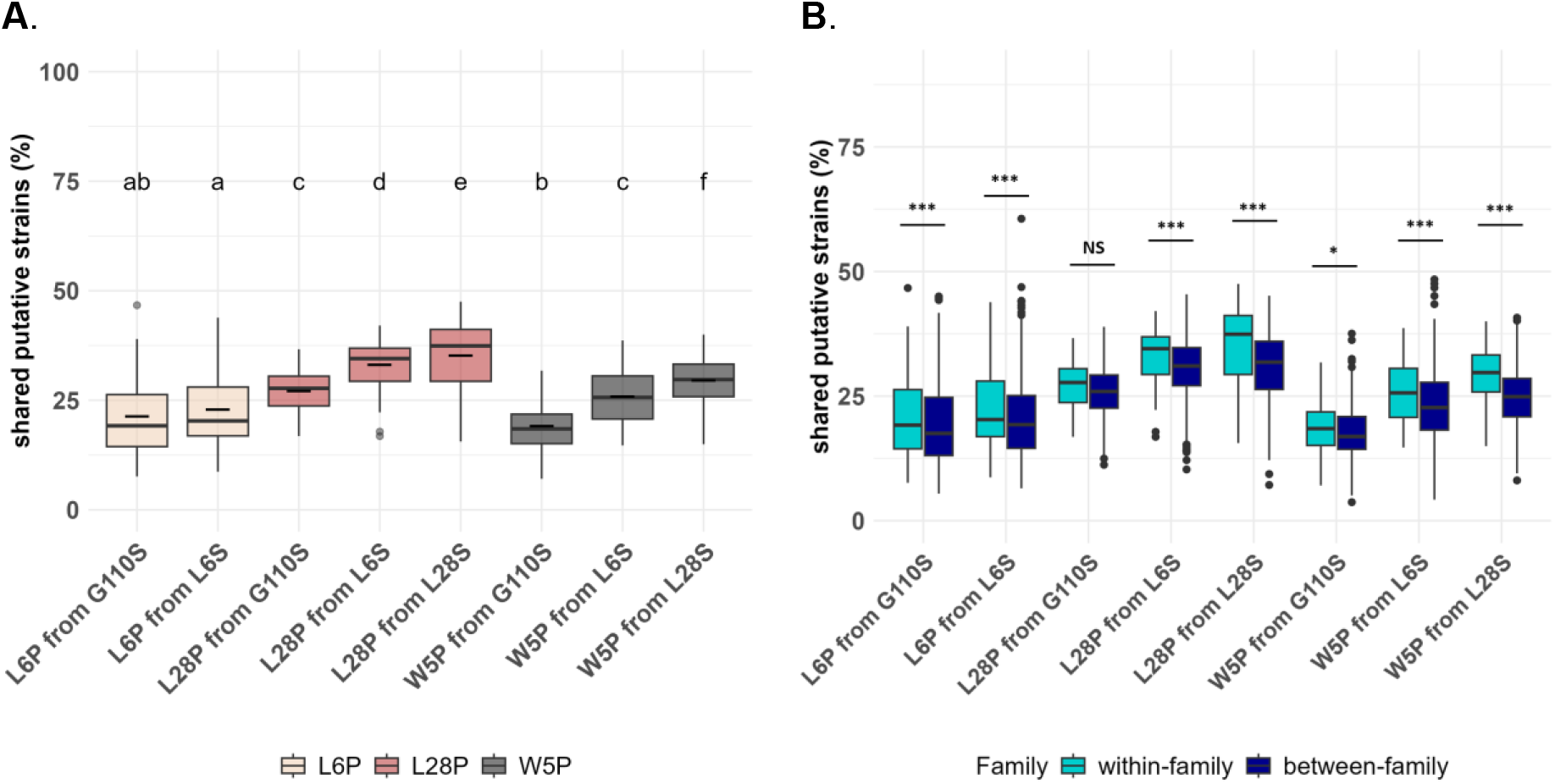
Proportion of putative strains shared between piglets and their mother at different stages. A. The beige boxplot corresponds to piglets at 6 days of lactation, the red boxplot to piglets at 28 days of lactation, and the grey boxplot to piglets at 5 days post-weaning. B. PS shared between related and unrelated sow-piglet are shown in cyan and dark blue respectively. L6P, L28P, and W5P correspond to piglet fecal samples collected at 6 and 28 days of suckling and 5 days after weaning, respectively; G110S, L6S, and L28S correspond to sow fecal samples collected at the end of gestation (day 110), at the beginning of lactation (day 6), and at the end of lactation (day 28, weaning), respectively. Different letters indicate significant differences between means (p < 0.05). n=17 sows, n= 51 piglets and 3 piglets per sow. NS > 0.05; * p < 0.05; *** p < 0.001.

Focusing on PS transmitted by the sow at the end of lactation *i*.*e*. shared with her piglets either at the end of lactation or 5 days after weaning, only one PS affiliated with *Limosilactobacillus reuteri* was consistently transmitted from the sow to all her piglets across all families (Figure 7A). Among the 50 most frequently transmitted strains, most were detected at both stages (before and after weaning), except for six PS. These six PS were preferentially transmitted at the end of lactation and were no longer detected after weaning. They were affiliated with *Romboutsia ilealis, Terrisporobacter petrolearius* (two PS) (Figure 7B), *Terrisporobacter mayombei, Turicibacter bilis*, and *Clostridium saudiense*. Interestingly, the 50 most frequently transmitted PS were not systematically the most abundant PS in sows at 28 days of lactation (R =0.15, p = 0.3 Figure 7C).

**Figure 7:**
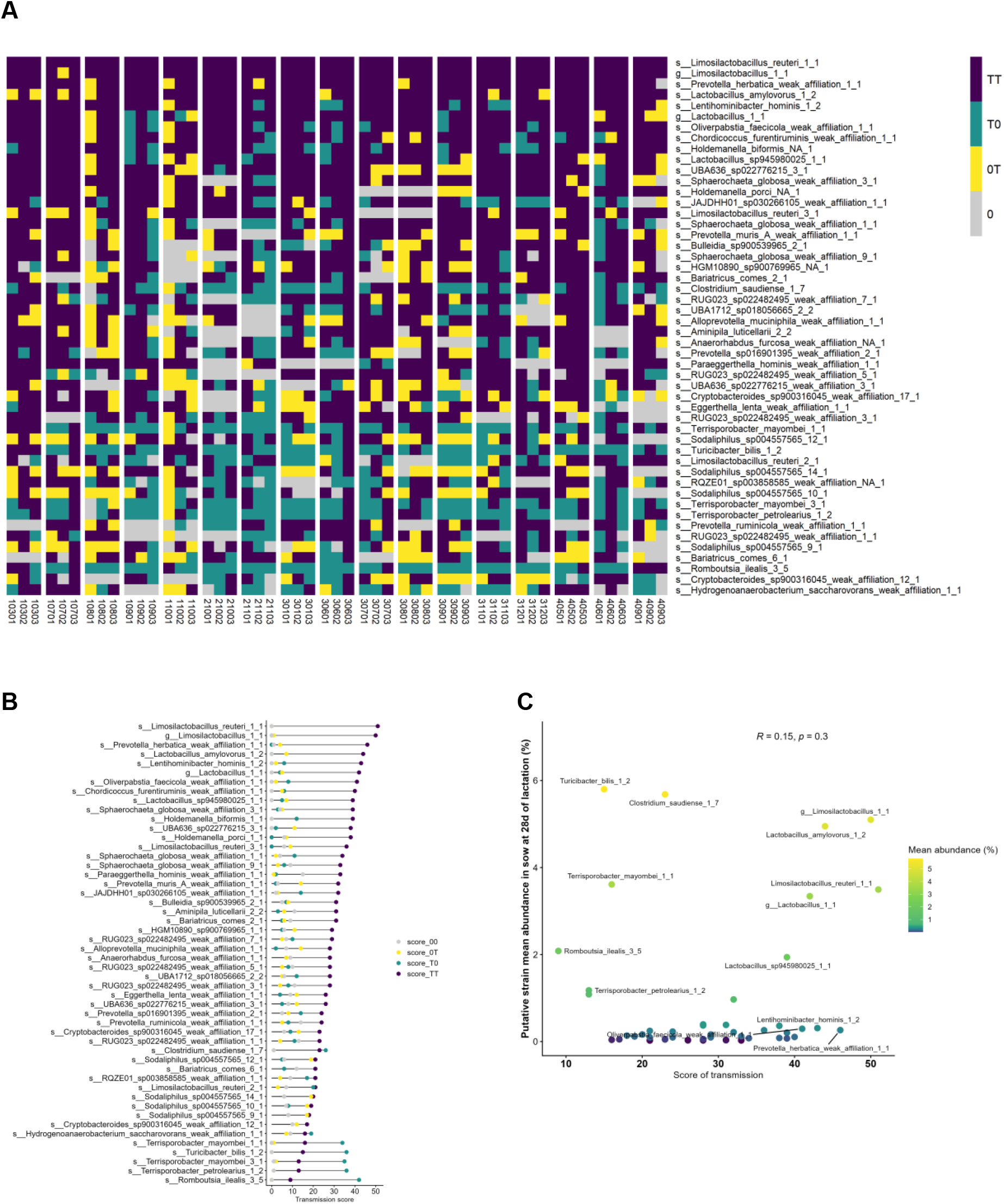
Transmitted putative strains (PS) from 28-day of lactation sows to 28-days-old suckling piglets and 5 days post-weaning piglets. A. Heatmap showing 50 transmitted PS to piglets with the highest transmission score. The rows correspond to the PS with taxonomic affiliations at the species (s_) or genus (g_) level. s_Limosilactobacillus_amylovorus_1_2 refers to PS 1 clustering two ASVs affiliated to *Limosilactobacillus amylovorus*. Weak_affilation refers to a BLAST identity lower than 95%. The columns represent the 51 piglets, grouped according by family (17 litters). Colors indicate PS transmission across stages: purple (TT), transmitted PS detected in piglets at both 28 days and 5 days post-weaning; green (T0), transmitted PS detected only in 28-day-old suckling piglets; yellow (0T), PS detected only in 5-day post-weaning piglets; grey (00) PS not detected at any of the stages. B. Lollipop plot showing the transmission score, defined as the sum of PS presence across stages: TT (detected in both stages), T0 (detected only in 28-day-old suckling piglets), 0T (detected only in 5-day post-weaning piglets), and 00 (not detected at any of the stages). C. Relationship between PS relative abundance in sow at 28 days of lactation and transmission score. Points are colored according to relative abundance (yellow: high; purple: low).

Overall, piglets shared significantly more PS with their sow during lactation than during late gestation. This pattern was evident at the end of lactation (L28P) and persisted after weaning (W5P). Furthermore, PS transmission was highest at the end of lactation, suggesting that this period represents a key window for sow-to-piglet microbiota transfer.

### Persistence of transmitted sow-derived putative strains in piglets during lactation and after weaning

Finally we assessed the persistence of PS transmitted from sow to her piglets by calculating the proportion of PS shared between related sows and piglets that persisted throughout lactation or were still present 5 days after weaning (Figure 8 and Table 2).

**Table 2:** Maternal transmitted and persistant PS in piglets.

| Maternal origin | Persistence in piglets | Number of transmitted and persistent PS per piglet | Number of family, genus, species<sup>1</sup> |
| --- | --- | --- | --- |
| G110S or L6S | L6P - L28P | $43 \pm 25^b$ | 46, 118, 168 |
| G110S or L6S or L28S | L28P - W5P | $122 \pm 56^a$ | 54, 177, 286 |
| G110S or L6S | L6P - L28P - W5P | $21 \pm 13^c$ | 35, 80, 110 |

<sup>1</sup> Number of bacterial families, genera and species assigned to transmitted and persistent PS
a,b,c Mean with different letter differ at $p < 0.05$

**Figure 8:**
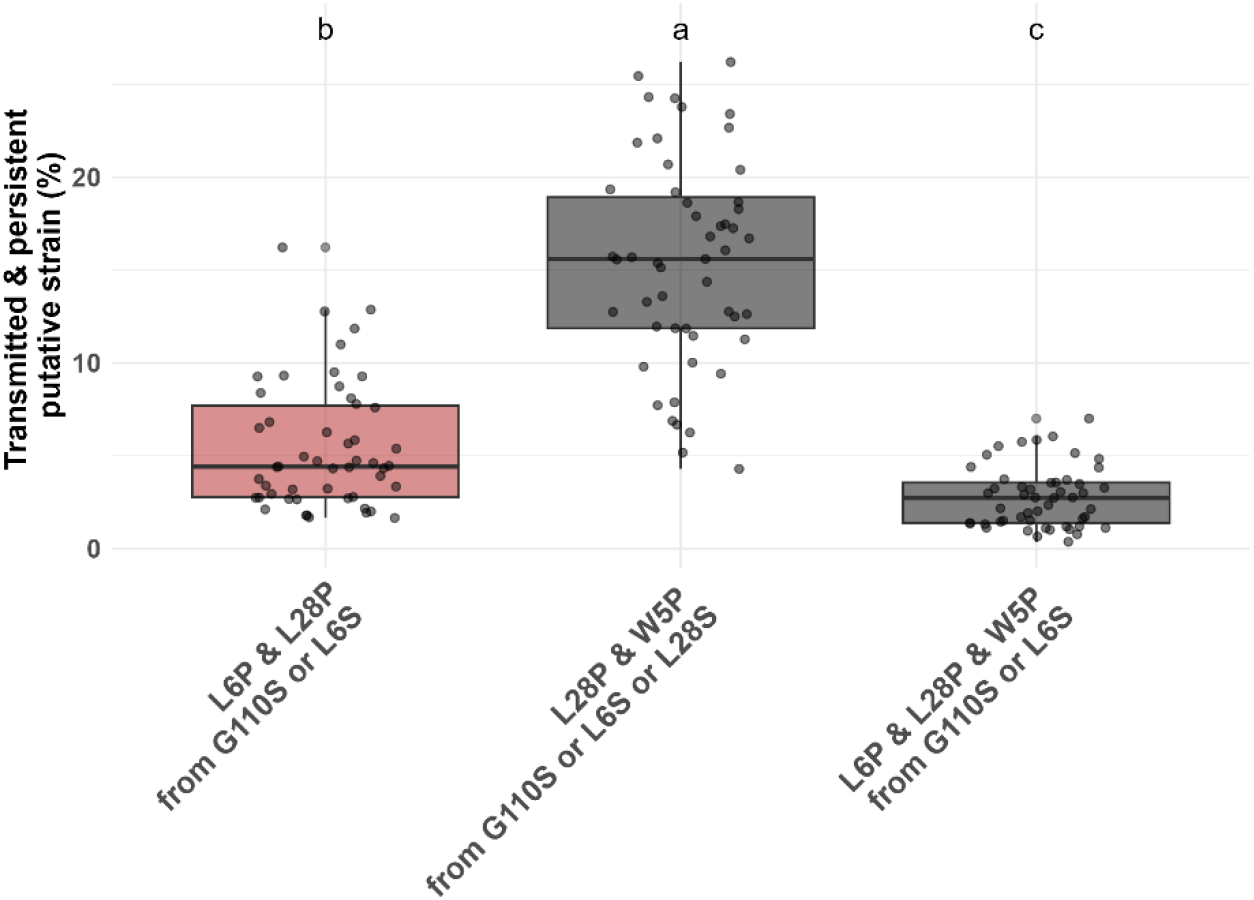
Proportion of putative strains shared between mothers and her piglets and persistant in piglets across different stages. Red boxplot refers to piglets at 28 days of lactation, and grey boxplot to piglets at 5 days post-weaning. L6P, L28P, and W5P correspond to piglet fecal samples collected at 6 and 28 days of suckling and 5 days after weaning, respectively; G110S, L6S, and L28S correspond to sow fecal samples collected at the end of gestation (day 110), at the beginning of lactation (day 6), and at the end of lactation (day 28, weaning), respectively. Different letters indicate significant differences between means (p < 0.05).

#### PS transmitted to piglets and persistent from day 6 to day 28 of lactation

PS transmitted to piglets and persistent during lactation represented 5.5 ± 3.4 % (Figure 8) i.e. 43 PS on average per piglets (Table 2). These PS belong to 46, 118 and 168 distincts bacterial families, genus and species respectively. During this period only Lactobacillaceae family with *Limosillactobacillus mucosae* and *Limosilactobacillus reuteri* species, Eggerthellaceae family with *Paraeggerthella hominis* and Anaerovoracaceae family were transmitted and persitent across all piglet families (Supplementary Figure S5, https://forge.inrae.fr/geraldine.pascal/sow-to-piglet-data-article).

#### PS transmitted to piglets and persistent from day 28 of lactation until 5 days after weaning

PS transmitted to piglets and persistent around weaning represented 15.4 ± 5.6 % (Figure 8). On average, 122 PS are shared from the sow between the end of lactation and remained detectable in her piglets after weaning (Table 2). These 122 PS were distributed across 54 bacterial families, 177 genera, and 286 species. PS that were both transmitted and persistent in at least one piglet across all families were restricted to 15 bacterial families (Supplementary Figure S6, https://forge.inrae.fr/geraldine.pascal/sow-to-piglet-data-article). Notably, some PS detected 5 days post-weaning appeared to be exclusively established at 28 days of age (Figure 9B). These included bacteria affiliated with *Prevotella* (e.g., *P. herbatica* (2 PS), *P. muris, P. ruminicola*), *Sphaerochaeta globosa* (4 PS), *Bariatricus comes* (2 PS), *Chordicoccus furentiruminis* although most of these affiliations had sequence identity scores below 95%.

**Figure 9:**
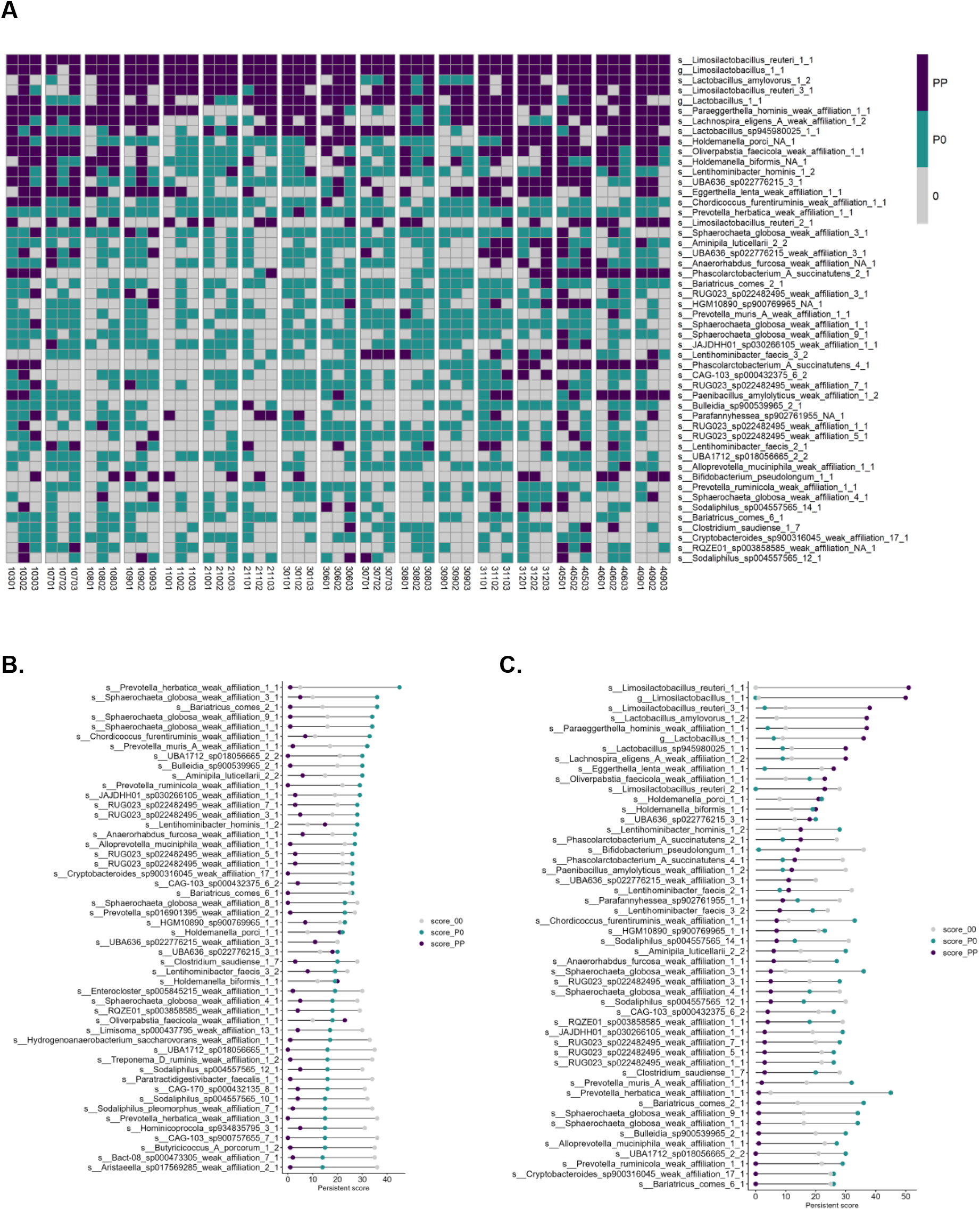
Transmitted and persistent putative strains (PS) in 5 days post-weaning piglets (top 50). A. Heatmap showing transmitted and persistent PS to piglets. The rows correspond to the PS with taxonomic affiliations at the species (s_) or genus (g_) level. s_Limosilactobacillus_amylovorus_1_2 refers to PS 1 clustering 2 ASVs affiliated to *Limosilactobacillus amylovorus*. Weak_affilation refers to a BLAST identity lower than 95%. The columns represent the 51 piglets, grouped according by family (17 litters). Colors indicate PS persitence across stages: purple (PP), transmitted PS detected in 5 days post-weaning piglets both from 6 days and from 28 days of lactation; green (P0), PS detected only in 28-day-old suckling piglets and in 5 days post-weaning piglets; grey (00) PS not detected at any of the stages. B and C. Lollipop plot showing the transmission score, defined as the sum of PS presence across stages: PP (detected in both stages), P0 (detected only in 28-day-old suckling piglets), and 00 (not detected at any of the stages). With ordering according to P0 (B panel) or PP score (C panel).

#### PS transmitted to piglets and persistent from day 6 of lactation to 5 days after weaning

The number of PS transmitted to piglets and persistent throughout all the three studied periods was the lowest (2.7 ± 1.6 % Figure 8). As expected considering persitence of PS within piglets, transmitted and persistent PS from day 6 of lactation until 5 days after weaning was limited to 21 PS on average. These 21 PS spanned 51 bacterial families, 80 genera, and 110 species. However, PS that were transmitted and persistent in at least one piglet in all family were restricted *Limosilactobacillus* genus (2 PS) including *L. reuteri* (Figure 9A & 9C & Supplementary Figure S7, https://forge.inrae.fr/geraldine.pascal/sow-to-piglet-data-article). PS affiliated with *Lactobacillus amylovorus* and *Paraeggerthella hominis* (affiliation score < 95%) was persistent across all litters except one or two respectively.

Altogether, persistent PS detected after weaning were predominantly transmitted at the end of lactation, supporting the idea that this period represents a critical window for the stable establishment of maternally derived strains in the piglet gut.

## Discussion

In pig production, weaning is a critical period for piglets, often associated with stress and digestive disorders (Lallès et al., 2007), which are commonly treated with antibiotics and may contribute to antimicrobial resistance (Paul et al., 2023). Early-life gut microbiota plays an important role in host health (Nowland et al., 2022). It is therefore hypothesised that controlling its establishment, particularly through the maternal transmission, could represent a promising strategy to better manage digestive disorders and reduce antibiotic use. Despite its importance, the proportion of microbiota transmitted from the sow to her piglets remains poorly characterized (Chen, Mi, et al., 2018; Liu et al., 2019; Tancredi et al., 2025). This study therefore aimed to better understand maternal microbiota transmission and persistance after weaning.

### Definition of putative strains from full-length amplicon data

To address this question, full-length 16S-ITS-23S amplicons were generated to achieve the highest possible taxonomic resolution. Because this resolution sometimes separated ribosomal operon copies into distinct ASVs, a secondary clustering step was performed: ASVs sharing the same species-level taxonomic affiliation and showing correlated abundance patterns were grouped together. These clusters, which regroup ASVs from the same genomes, were defined as putative strains (PS). The resulting reduction of dataset complexity remained limited: more than two-thirds of ASVs were not merged with any others, either because their abundances were too low to reliably assess correlations or because ribosomal operon copies in their originating genome were already similar enough to have been merged during ASV inference. Interestingly, alpha- and beta-diversity patterns based on PS were consistent with the typical dynamics of fecal bacterial communities previously described in sows and piglets using short read amplicon (Le Floc’h et al., 2022; Tancredi et al., 2025).

To better understand maternal microbiota transmission and persistance of PS after weaning in their offspring we followed a three-step analysis. First we evaluated PS persistence within both partners (sows and piglets), then we evaluated transmission of PS from sow to her offsprings and finaly we combined the latter to evaluated the transmitted persistent PS.

### PS persistence in sows

In sows, we observed that nearly half of the putative strains persisted between consecutive physiological stages, from late gestation to late lactation. However, persistence decreased when considering the entire period, indicating a gradual restructuring of the maternal gut microbiota over time. During this period, sows undergo major physiological changes associated with parturition and the onset of lactation, along with dietary adjustments to meet the nutritional demands of lactation. Previous studies have described microbiota dynamics associated with diet and physiological state (Leblois et al., 2018). For instance, a previous study reported a parturition-related transient decrease, followed by recovery within 5 days postpartum, of several taxa including *Limosilactobacillus reuteri, Lactobacillus amylovorus, Lactobacillus johnsonii*, and *Limosilactobacillus mucosae*, as well as members of the *Clostridium leptum* and *Clostridium coccoides* groups (Paßlack et al., 2015). Furthermore at the end of gestation, *Lactobacillus amylovorus* and *Limosilactobacillus reuteri* were differentially abundant according to sow parity, with significant differences observed between nulliparous and high-parity sows (Berry et al., 2021). In the present study, the dynamics of persistent PS from late gestation to the end of lactation were found to be PS-specific. *Limosilactobacillus mucosae* decreased after parturition and increased again at the end of lactation, whereas *Limosilactobacillus reuteri* exhibited the opposite pattern. Persistent PS, affiliated with *Lactobacillus amylovorus*, sharply decreased after parturition and remained stable thereafter. These results highlight differential responses of microbiota members to physiological and dietary changes in sows, and underscore the importance of high-resolution approaches to capture fine-scale microbial dynamics that may be overlooked using lower-resolution methods.

### PS persistence in piglets

In contrast, the proportion of persistent PS between the beginning and the end of lactation was lower in piglets than in sows. This low persistence may reflect the major remodeling of the gut microbiota with a strong ecological species succession occurring during early-life during microbiota assembly (Chen, Xu, et al., 2018; Luo et al., 2022). Initial colonizers are progressively replaced as the gut environment matures, driven by host development and dietary transitions. In piglets, the highest persistence was observed between the end of lactation and 5 days after weaning as previously observed (Tancredi et al., 2025). Although still lower than in sows, this increased persistence suggests the emergence of a more stable microbiota driven by the introduction of solid food which occured in our study at 21 days of lactation. Persistent PS detected across piglets all along the sampling process, from 6 days of lactation on, were affiliated at the species level with *Limosilactobacillus reuteri (3 PS)* and *Lactobacillus amylovorus* followed with *Holdemanella porci and H. biformis, Lentihominibacter hominis* and *Dorea formicigenerans*. Their persistence was not only linked to their abundance but likely reflects their high capacity of adaptation to both diet changing and host gut maturation. For instance, aerotolerant anaerobes such as *Limosilactobacillus reuteri* and *Lactobacillus amylovorus*, commonly reported in the pig gut (Højberg et al., 2005), are well adapted to the early-life gut, with traits including acid and bile resistance, epithelial adhesion, and antimicrobial activity that support stable colonization (Kim et al., 2007; Yang et al., 2020). Notably, *L. reuteri* has been linked to microbiota maturation in piglets (Wang et al., 2022). In parallel, strictly anaerobic taxa such as *Holdemanella* spp. (De Maesschalck et al., 2014; Wylensek et al., 2020) and *D. formicigenerans* (Holdeman & Moore, 1974) contribute to carbohydrate fermentation and short-chain fatty acid production, indicating early establishment of anaerobic functions. However, such persistence may also reflect repeated re-inoculation events, driven by continuous maternal transmission, thereby supporting a sustained transmission of specific strains during lactation.

### Sow-to-piglet transmission: a stage-dependent dynamic

Our analysis confirms significant maternal transmission of gut microbiota, as evidenced by the higher proportion of shared PS between sows and their own piglets compared to unrelated pairs. However, this transmission was strongly stage-dependent. The lowest proportion was observed at the beginning of lactation, suggesting that, at this stage, the neonatal gut microbiota is still largely shaped by other colonization sources and that the intestinal environment may not yet be favorable for the establishment of maternal strains. In line with this, previous studies have shown that early microbial communities in piglets are more similar to environmental sources (e.g., floor, milk, and nipple surfaces) than to maternal fecal microbiota, with initial colonizers originating from multiple maternal and environmental reservoirs (Chen, Xu, et al., 2018). In contrast, the proportion of shared PS derived from the sow was highest at the end of lactation, both before and after weaning. These results are consistent with previous findings showing increasing similarity between sow and piglet fecal microbiota over the course of lactation (Chen, Xu, et al., 2018), and support the idea that late lactation represents a key window for sow-to-piglet microbiota transmission. However, these results differ from those of Tancredi et al., (2025), who reported that the contribution of the maternal microbiota to piglets is highest at the beginning of lactation, decreases markedly toward the end of lactation, and then increases again within 5 to 10 days after weaning. In our study, five days after weaning, the proportion of PS derived from the sow at L28 that were shared with piglets decreased compared with the end of lactation. This pattern is consistent with the cessation of direct contact between sows and piglets after weaning and suggests that maternal-associated PS may progressively be lost or become less abundant once maternal exposure ceases. The increase in maternal contribution reported by Tancredi et al. (2025) after weaning, particularly in the absence of further direct contact with the sow, might reflect the subsequent proliferation of bacteria acquired from the sow earlier in life that were previously present at non-detectable abundance. Differences in the approaches used to infer maternal contribution may also account for the contrasting patterns observed between studies. Furthermore, differences in experimental design, including the number of sows and piglets involved, the resolution of bacterial tracking (*i*.*e*., species *vs*. genus), as well as environmental exposure and husbandry conditions (*i*.*e*., antibiotic treatment, housing systems, and hygiene levels), may all substantially influence estimates of maternal microbial transmission. Focusing on strains shared with the sows at the end of lactation, only one PS affiliated with *Limosilactobacillus reuteri* was consistently transmitted and remained detectable 5 days after weaning in all piglets from all families. Notably, the PS transmission success seemed not to be correlated with its abudance in sows, in agreement with previous findings comparing sows at 114 days of gestation and piglets at 10 days of age (Berry et al., 2021). Several PS, including those affiliated with *Romboutsia ilealis, Terrisporobacter petrolearius, Terrisporobacter mayombei, Turicibacter bilis*, and *Clostridium saudiense*, were transmitted during late lactation but did not persist after weaning. This lack of persistence may reflect the ecological disturbance associated with weaning, including dietary changes, environmental stress, and gut physiological remodeling, which can impose selective pressures unfavorable to certain maternal strains. Consequently, only a subset of transmitted bacteria may successfully establish and persist in the post-weaning gut environment.

### Transmitted PS persisting after weaning

Assuming that maternal transmission of beneficial bacteria contributes to shaping a favorable gut microbiota in piglets we identified PS that were both shared with the sow at different stages and persistent in at least one piglet within the litter after weaning. Not surprisingly, the number of such persistent shared PS was further reduced. The persistence of transmitted ASVs from sow to offspring after weaning was previously demonstrated using V3-V4 metabarcoding (Tancredi et al., 2025), but never at the taxonomic resolution achieved with full-length 16S-ITS-23S sequencing. The highest number of sow-derived persitent PS in piglet was observed for PS transmitted at the end of lactation. From an ecological perspective, these maternally shared PS likely benefit from a sufficiently similar niche both at the end of lactation and in the post-weaning gut environment, which is largely driven by the transition to solid feed. Persistent transmitted PS were mainly affiliated with *Prevotella* species (including PS near *P. herbatica* and *P. muris*), as well as *Bariatricus comes* and S*phaerochaeta globosa. Prevotella* is a dominant genus in the porcine gut (Larzul et al., 2024; Tancredi et al., 2025), known for its ability to degrade complex plant polysaccharides (Accetto & Avguštin, 2019). *Bariatricus comes (*formerly *Coprococcus comes)*, an obligately anaerobic member of the Lachnospiraceae displays a comparatively narrow carbohydrate utilisation profile, growing well primarily on glucose, which it ferments into lactate, butyrate, and acetate (Notting et al., 2023). Regarding *Sphaerochaeta globosa*, this obligately anaerobic spirochete ferments a range of mono-, di-, and polysaccharides into acetate, formate, and ethanol (Ritalahti et al., 2012), and its genome further encodes a moderately diverse repertoire of carbohydrate-active enzymes targeting plant-derived polysaccharides such as pectin, consistent with weak but stable growth observed on this substrate (Troshina et al., 2024)

Conversely, PS affiliated with *Limosilactobacillus* (including *Limosilactobacillus reuteri*), *Lactobacillus amylovorus*, and *Paraeggerthella hominis* were detected in the sow and persisted in piglets from day 6 of lactation until 5 days after weaning. The persistence of these strains across this extended period highlights their remarkable ability to adapt to highly distinct gut environments, ranging from the maternal intestine to that of suckling piglets and through the dietary transition associated with weaning. However, the number of such early-transmitted and persistent PS remained limited. This suggests that only a restricted subset of bacteria possesses the capacity to tolerate major dietary and environmental shifts, possibly through metabolic flexibility or niche specialization.

In conclusion, this study provides a high-resolution view of maternal microbiota transmission in pigs and its contribution to the establishment of the post-weaning gut microbiota. By combining species-level amplicon sequencing with a strain-oriented approach, we show that while maternal transmission occurs throughout lactation, only a limited subset of strains successfully persists after weaning. This persistence is strongly dependent on the timing of transmission, with late lactation emerging as the main window for the transfer of strains capable of establishing in the post-weaning gut. In contrast, strains transmitted earlier in life are largely transient, despite continuous exposure of piglets to their mother. However, the maternal microbiota present at the end of gestation and early lactation may still play an important role in early-life gut development, particularly in the control of neonatal diarrhea. Our results further indicate that persistence is not driven by abundance in the sow but rather by the ecological fitness of specific taxa to withstand major environmental transitions, particularly the shift to solid feeding. Persistent strains were mainly affiliated with carbohydrate-degrading and anaerobic taxa, highlighting the role of dietary-driven ecological selection in shaping microbiota establishment. Altogether, these findings improve our understanding of maternal microbial transmission dynamics in pigs and suggest that targeting the sow microbiota during late lactation could represent a promising strategy to promote the establishment of beneficial bacteria in piglets and potentially mitigate post-weaning disorders.

## Acknowledgements

We are grateful to the genotoul bioinformatics platform Toulouse Occitanie (Bioinfo Genotoul, https://doi.org/10.15454/1.5572369328961167E12) for providing computing and storage resources. We thank Laurent Cauquil for depositing the 16S-ITS-23S sequences in the European Nucleotide Archive (ENA).

## Data, scripts, code, and supplementary information availability

Animal metadata are available online: TableS1 Animal_metadata_supplemental260206.xlsx https://doi.org/10.57745/3ZJXTI

Scripts, and are available online: https://forge.inrae.fr/geraldine.pascal/sow-to-piglet-data-article

Supplementary figures are available online https://forge.inrae.fr/geraldine.pascal/sow-to-piglet-data-article

Sequences were deposited in European Nucleotide Archive (ENA): accession number is PRJEB113713.

