## Supplemental figures for "Late lactation represents the main window for sow-to-piglet transmission of persistent gut strains"

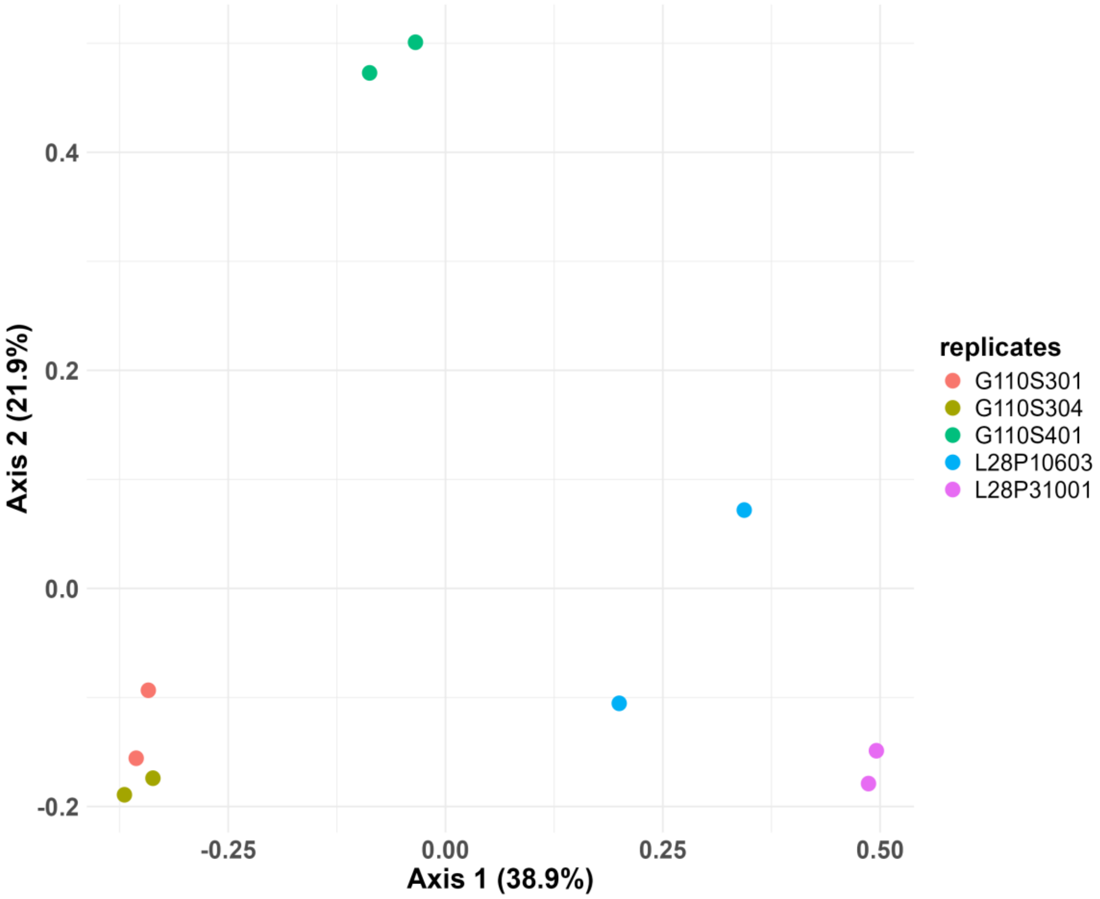


**Figure S1:** Principal Coordinates Analysis (PCoA) based on Bray-Curtis distance of putative strain composition from five fecal samples, each extracted and sequenced in duplicate, to assess the repeatability of the sequencing results. L6P, L28P, and W5P correspond to piglet fecal samples collected at 6 and 28 days of suckling and 5 days after weaning, respectively; Sample starting with G110S correspond to sow fecal samples collected at the end of gestation (day 110); sample starting with L28P correspond to piglet fecal samples collected at 28 days of suckling.

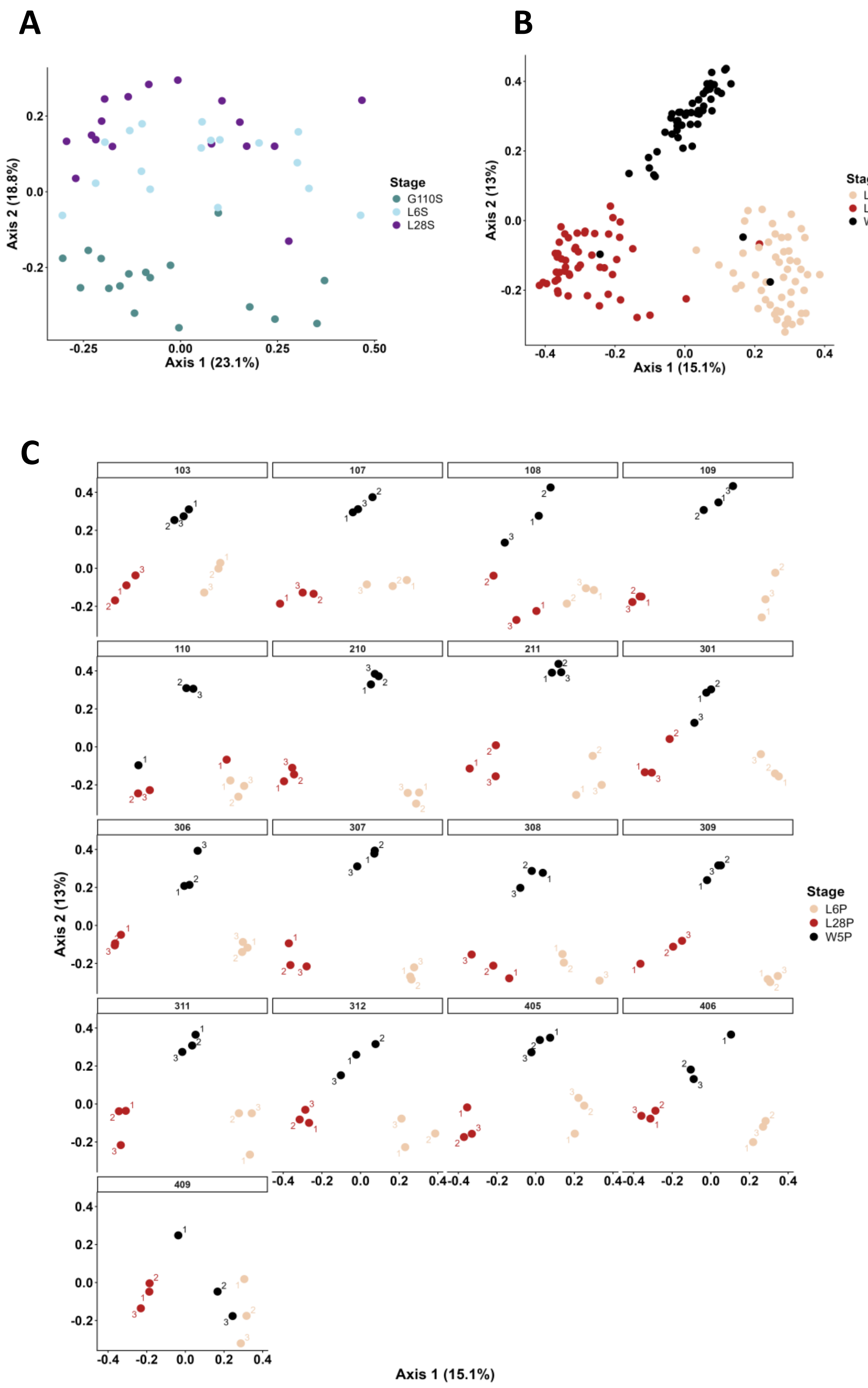


**Figure S2:** PCoA based on Bray-Curtis distance of putative strain composition in piglet and sow fecal samples. A. PCoA calculated on sow fecal samples (n = 17 per sow stage). B. PCoA calculated on piglet fecal samples (n = 51 per stage). C. Same PCoA as in B, split per litter (n = 17 litters, 3 piglets per litter, followed across three stages). G110S, L6S, and L28S correspond to sow fecal samples collected at the end of gestation (day 110), at the beginning of lactation (day 6), and at the end of lactation (day 28, weaning), respectively; L6P, L28P, and W5P correspond to piglet fecal samples collected at 6 and 28 days of suckling and 5 days after weaning

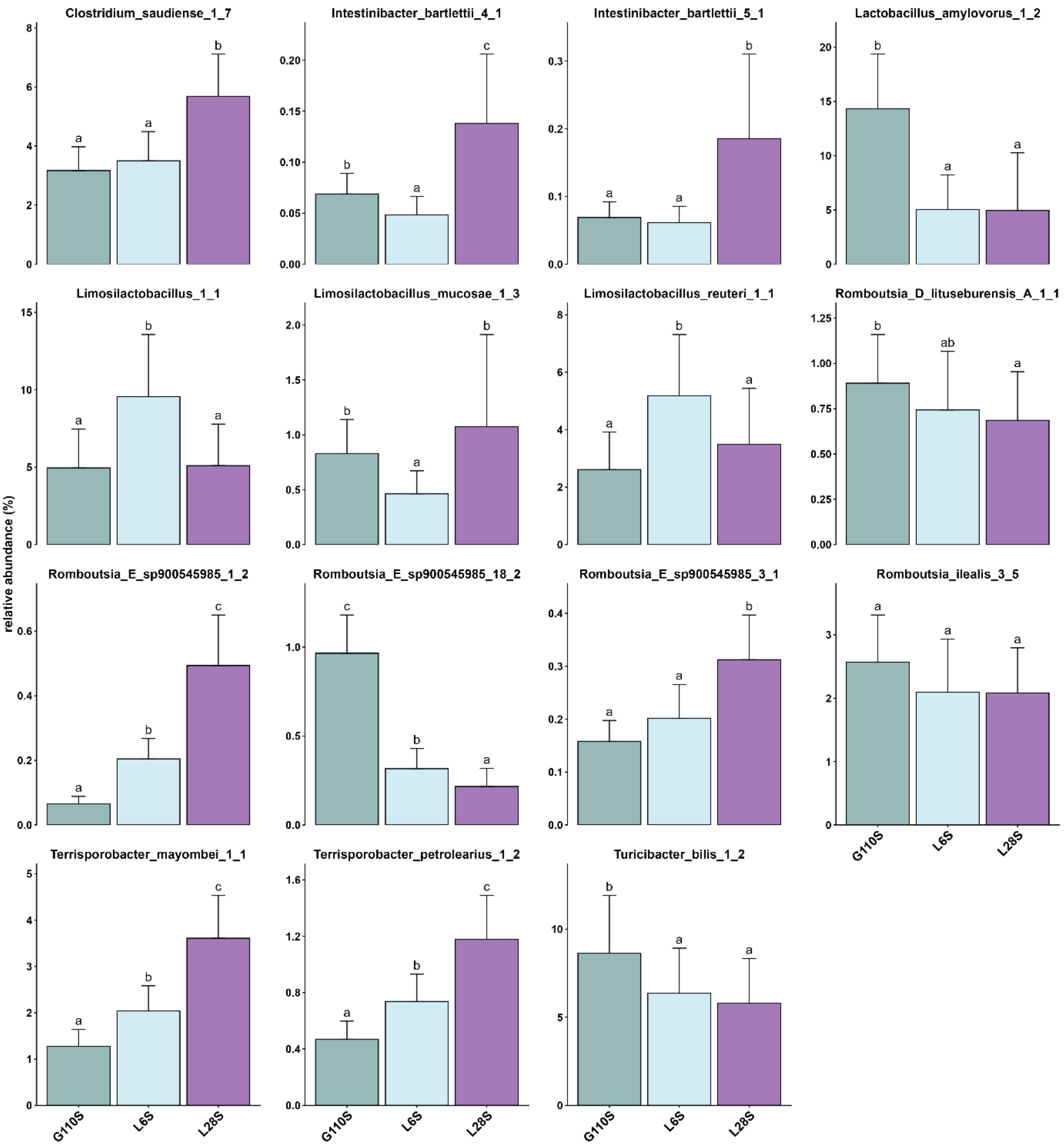


**Figure S3:** Relative abundance of the 15 persistent putative strains present in all sows from the end of gestation throughout lactation. G110S, L6S, and L28S correspond to sow fecal samples collected at the end of gestation (day 110), at the beginning of lactation (day 6), and at the end of lactation (day 28, weaning), respectively (n = 17 sows). Clostridium_saudiense_1_7 refers to PS 1 clustering seven ASVs affiliated to *Clostridium saudiense.* Different letters indicate a significant difference between means ($p < 0.05$)

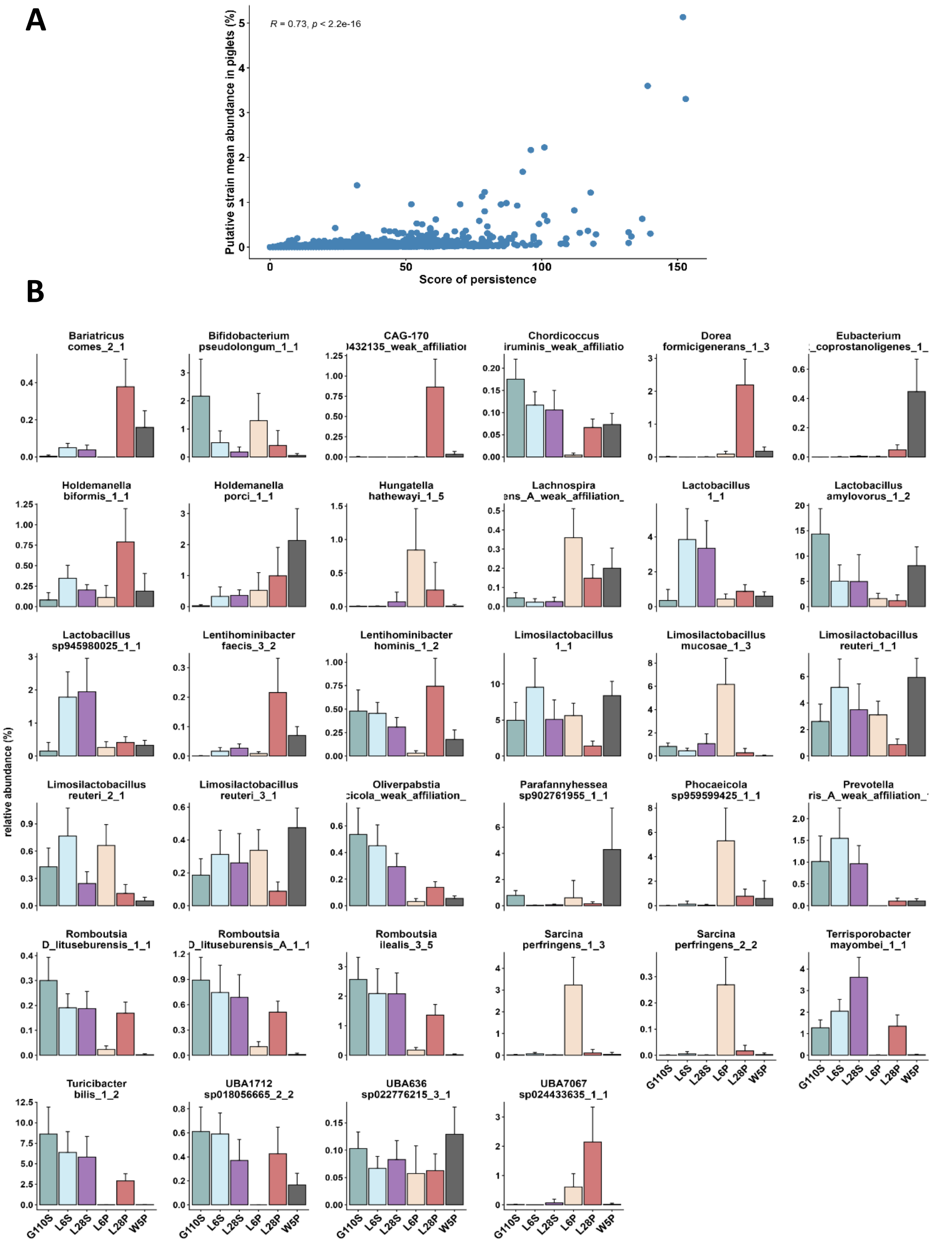


**Figure S4:** Putative strain (PS) persistence in piglet fecal samples A. Spearman correlation between PS persistence score and mean abundance in piglets B. relative abundance in sows and piglets of the 34 persistent PS detected in all piglets from the 17 litters. Bariatricus_comes_2_1 refers to PS 2 clustering one ASV affiliated to *Bariatricus comes*. G110S, L6S, and L28S correspond to sow fecal samples collected at the end of gestation (day 110), at the beginning of lactation (day 6), and at the end of lactation (day 28, weaning), respectively (n = 17 sows). L6P, L28P, and W5P correspond to piglet fecal samples collected at 6 and 28 days of suckling and 5 days after weaning (n = 51 piglets).

A.

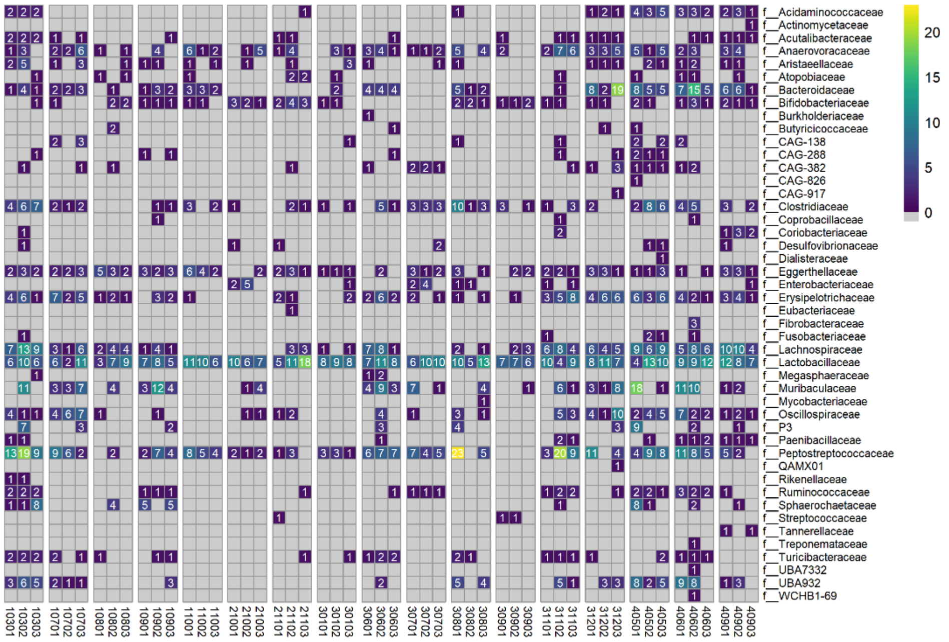
f__Acidaminococcaceae
f__Actinomycetaceae
f__Acutalibacteraceae
f__Anaerovoracaceae
f__Aristaeellaceae
f__Atopobiaceae
f__Bacteroidaceae
f__Bifidobacteriaceae
f__Burkholderiaceae
f__Butyricicoccaceae
f__CAG-138
f__CAG-288
f__CAG-382
f__CAG-826
f__CAG-917
f__Clostridiaceae
f__Coprobacillaceae
f__Coriobacteriaceae
f__Desulfovibrionaceae
f__Dialisteraceae
f__Eggerthellaceae
f__Enterobacteriaceae
f__Erysipelotrichaceae
f__Eubacteriaceae
f__Fibrobacteraceae
f__Fusobacteriaceae
f__Lachnospiraceae
f__Lactobacillaceae
f__Megasphaeraceae
f__Muribaculaceae
f__Mycobacteriaceae
f__Oscillospiraceae
f__P3
f__Paenibacillaceae
f__Peptostreptococcaceae
f__QAMX01
f__Rikenellaceae
f__Ruminococcaceae
f__Sphaerochaetaceae
f__Streptococcaceae
f__Tannerellaceae
f__Treponemataceae
f__Turicibacteraceae
f__UBA7332
f__UBA932
f__WCHB1-69
20
15
10
5
0
10301 10302 10303 10701 10702 10703 10801 10802 10803 10901 10902 10903 11001 11002 11003 21001 21002 21003 21101 21102 21103 30101 30102 30103 30601 30602 30603 30701 30702 30703 30801 30802 30803 30901 30902 30903 31101 31102 31103 31201 31202 31203 40501 40502 40503 40601 40602 40603 40901 40902 40903

B.

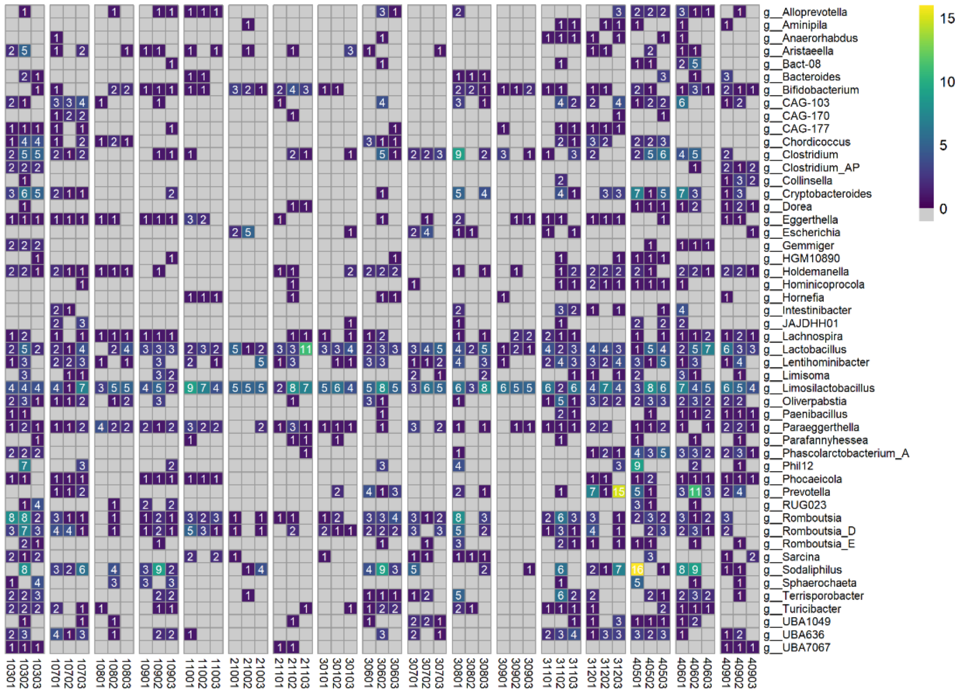
g__Alloprevotella
g__Aminipila
g__Anaerorhabdus
g__Aristaeella
g__Bact-08
g__Bacteroides
g__Bifidobacterium
g__CAG-103
g__CAG-170
g__CAG-177
g__Chordicoccus
g__Clostridium
g__Clostridium_AP
g__Collinsella
g__Cryptobacteroides
g__Dorea
g__Eggerthella
g__Escherichia
g__Gemmiger
g__HGM10890
g__Holdemanella
g__Hominicoprocola
g__Hornefia
g__Intestinibacter
g__JAJDHH01
g__Lachnospira
g__Lactobacillus
g__Lentihominibacter
g__Limisoma
g__Limosilactobacillus
g__Oliverpabstia
g__Paenibacillus
g__Paraeggerthella
g__Parafannyhessea
g__Phascolarctobacterium_A
g__Phil12
g__Phocaeicola
g__Prevotella
g__RUG023
g__Romboutsia
g__Romboutsia_D
g__Romboutsia_E
g__Sarcina
g__Sodaliphilus
g__Sphaerochaeta
g__Terrisporobacter
g__Turicibacter
g__UBA1049
g__UBA636
g__UBA7067
15
10
5
0
10301 10302 10303 10701 10702 10703 10801 10802 10803 10901 10902 10903 11001 11002 11003 21001 21002 21003 21101 21102 21103 30101 30102 30103 30601 30602 30603 30701 30702 30703 30801 30802 30803 30901 30902 30903 31101 31102 31103 31201 31202 31203 40501 40502 40503 40601 40602 40603 40901 40902 40903

**C.**

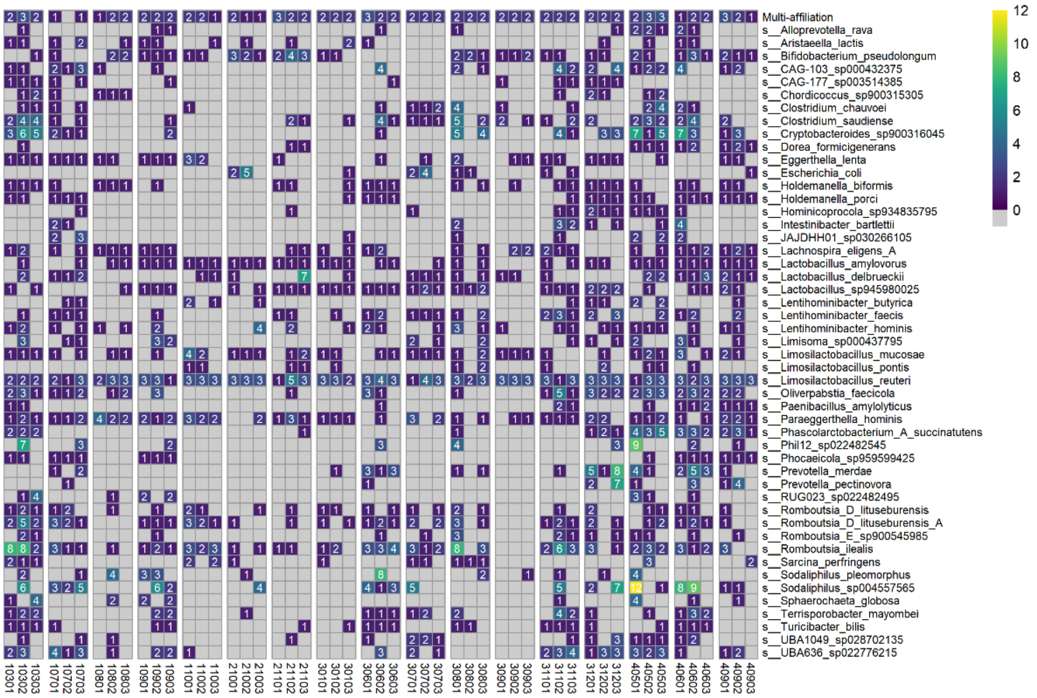


**Figure S5:** Number of G110- or L6-derived sow fecal transmitted and persistent putative strains in L6 and L28 piglet fecal samples. A. With affiliation at the family level. B. With affiliation at the genus and C. species level. For genus and species affiliation, only the top 50 taxa are presented. (G110: end of gestation, L6: 6 days of lactation, L28: 28 days of lactation)

A.

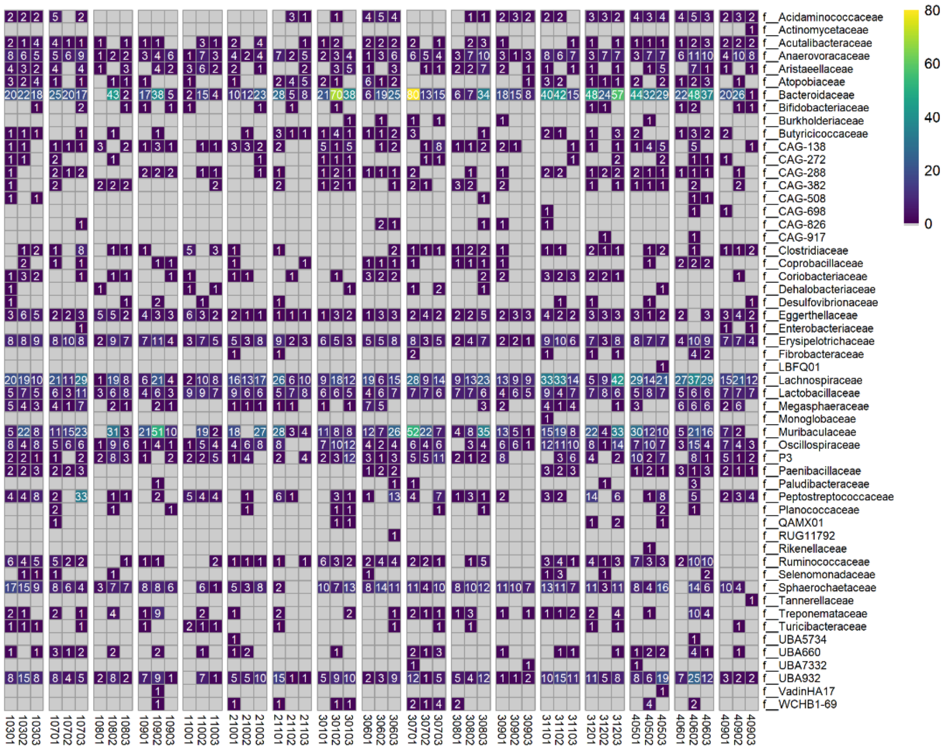
80
60
40
20
0
f__Acidaminococcaceae
f__Actinomycetaceae
f__Acutalibacteraceae
f__Anaerovoracaceae
f__Aristaeellaceae
f__Atopobiaceae
f__Bacteroidaceae
f__Bifidobacteriaceae
f__Burkholderiaceae
f__Butyricicoccaceae
f__CAG-138
f__CAG-272
f__CAG-288
f__CAG-382
f__CAG-508
f__CAG-698
f__CAG-826
f__CAG-917
f__Clostridiaceae
f__Coprobacillaceae
f__Coriobacteriaceae
f__Dehalobacteriaceae
f__Desulfovibrionaceae
f__Eggerthellaceae
f__Enterobacteriaceae
f__Erysipelotrichaceae
f__Fibrobacteraceae
f__LBFQ01
f__Lachnospiraceae
f__Lactobacillaceae
f__Megasphaeraceae
f__Monoglobaceae
f__Muribaculaceae
f__Oscillospiraceae
f__P3
f__Paenibacillaceae
f__Paludibacteraceae
f__Peptostreptococcaceae
f__Planococcaceae
f__QAMX01
f__RUG11792
f__Rikenellaceae
f__Ruminococcaceae
f__Selenomonadaceae
f__Sphaerochaetaceae
f__Tannerellaceae
f__Treponemataceae
f__Turicibacteraceae
f__UBA5734
f__UBA660
f__UBA7332
f__UBA932
f__VadinHA17
f__WCHB1-69
10301 10302 10303 10701 10702 10703 10801 10802 10803 10901 10902 10903 11001 11002 11003 21001 21002 21003 21101 21102 21103 30101 30102 30103 30601 30602 30603 30701 30702 30703 30801 30802 30803 30901 30902 30903 31101 31102 31103 31201 31202 31203 40501 40502 40503 40601 40602 40603 40901 40902 40903

B.

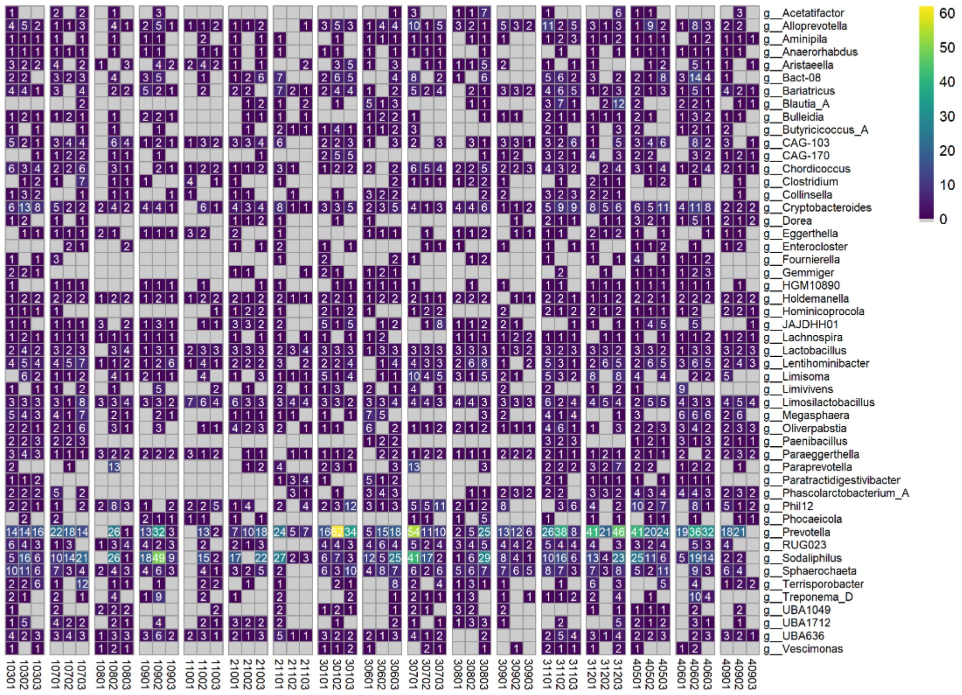
60
50
40
30
20
10
0
g__Acetatifactor
g__Alloprevotella
g__Aminipila
g__Anaerorhabdus
g__Aristaeella
g__Bact-08
g__Bariatricus
g__Blautia_A
g__Bulleidia
g__Butyricicoccus_A
g__CAG-103
g__CAG-170
g__Chordicoccus
g__Clostridium
g__Collinsella
g__Cryptobacteroides
g__Dorea
g__Eggerthella
g__Enterocloster
g__Fournierella
g__Gemmiger
g__HGM10890
g__Holdemanella
g__Hominicoprocola
g__JAJDHH01
g__Lachnospira
g__Lactobacillus
g__Lentihominibacter
g__Limisoma
g__Limivivens
g__Limosilactobacillus
g__Megasphaera
g__Oliverpabstia
g__Paenibacillus
g__Paraeggerthella
g__Paraprevotella
g__Paratractidigestivibacter
g__Phascolarctobacterium_A
g__Phil12
g__Phocaeicola
g__Prevotella
g__RUG023
g__Sodaliphilus
g__Sphaerochaeta
g__Terrisporobacter
g__Treponema_D
g__UBA1049
g__UBA1712
g__UBA636
g__Vescimonas
10301 10302 10303 10701 10702 10703 10801 10802 10803 10901 10902 10903 11001 11002 11003 21001 21002 21003 21101 21102 21103 30101 30102 30103 30601 30602 30603 30701 30702 30703 30801 30802 30803 30901 30902 30903 31101 31102 31103 31201 31202 31203 40501 40502 40503 40601 40602 40603 40901 40902 40903

C.

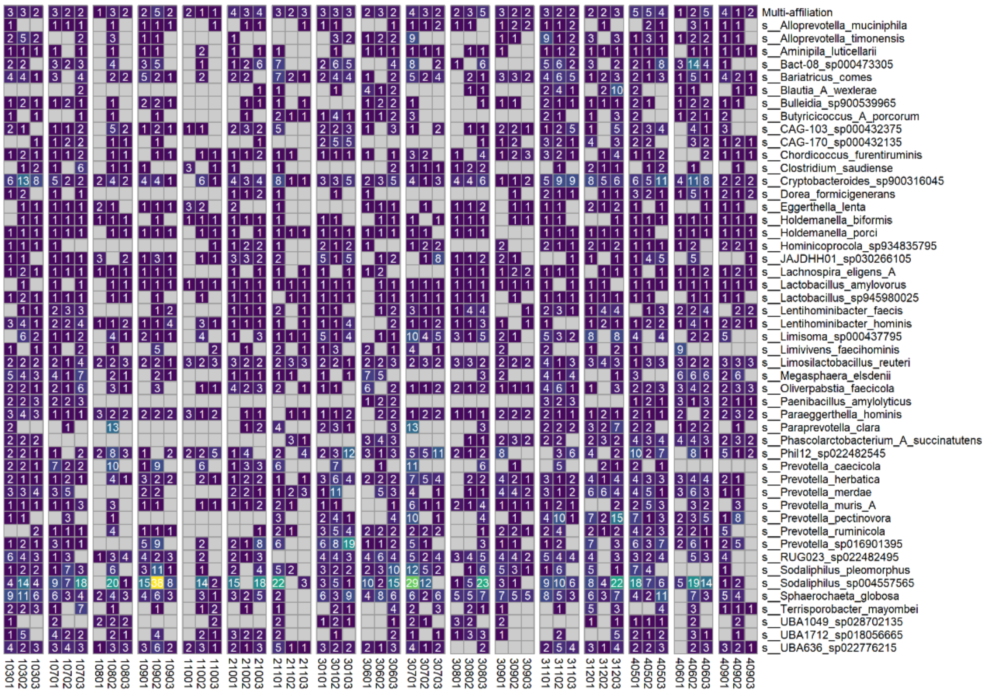


**Figure S6:** Number of G110- or L6- or L28-derived sow fecal transmitted and persistent putative strains in L28 and W5 piglet fecal samples. A. With affiliation at the family level. B. With affiliation at the genus and C. species level. For genus and species affiliation, only the top 50 taxa are presented. (G110: end of gestation, L6: 6 days of lactation, L28: 28 days of lactation, W5 : 5 days after weaning))

A.

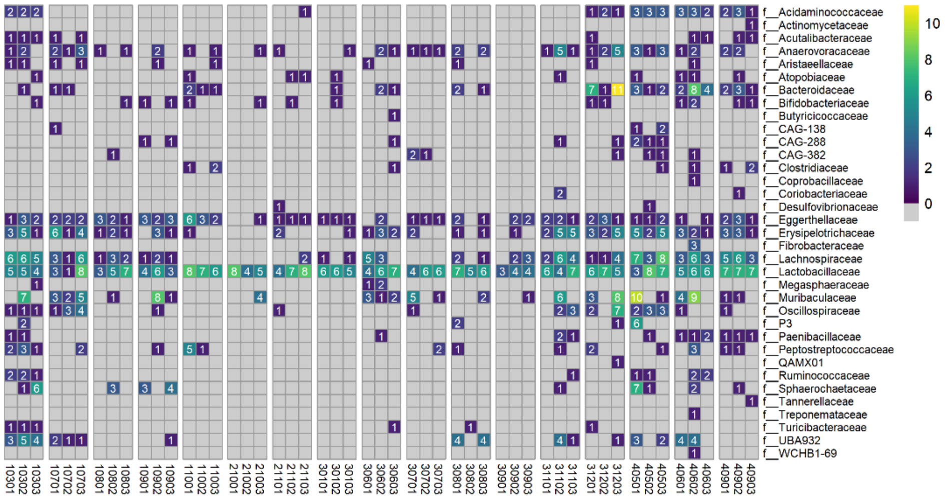
f__Acidaminococcaceae
f__Actinomycetaceae
f__Acutalibacteraceae
f__Anaerovoracaceae
f__Aristaeellaceae
f__Atopobiaceae
f__Bacteroidaceae
f__Bifidobacteriaceae
f__Butyricicoccaceae
f__CAG-138
f__CAG-288
f__CAG-382
f__Clostridiaceae
f__Coprobacillaceae
f__Coriobacteriaceae
f__Desulfovibrionaceae
f__Eggerthellaceae
f__Erysipelotrichaceae
f__Fibrobacteraceae
f__Lachnospiraceae
f__Lactobacillaceae
f__Megasphaeraceae
f__Muribaculaceae
f__Oscillospiraceae
f__P3
f__Paenibacillaceae
f__Peptostreptococcaceae
f__QAMX01
f__Ruminococcaceae
f__Sphaerochaetaceae
f__Tannerellaceae
f__Treponemataceae
f__Turicibacteraceae
f__UBA932
f__WCHB1-69
10
8
6
4
2
0
10301 10302 10303 10701 10702 10703 10801 10802 10803 10901 10902 10903 11001 11002 11003 21001 21002 21003 21101 21102 21103 30101 30102 30103 30601 30602 30603 30701 30702 30703 30801 30802 30803 30901 30902 30903 31101 31102 31103 31201 31202 31203 40501 40502 40503 40601 40602 40603 40901 40902 40903

B.

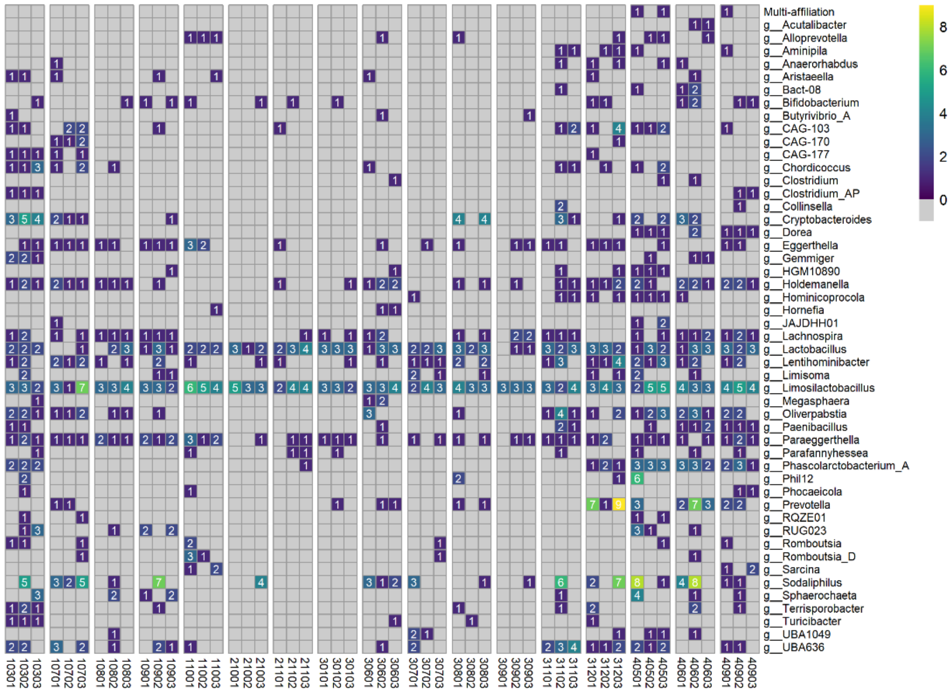
Multi-affiliation
g__Acutalibacter
g__Alloprevotella
g__Aminipila
g__Anaerorhabdus
g__Aristaeella
g__Bact-08
g__Bifidobacterium
g__Butyrivibrio_A
g__CAG-103
g__CAG-170
g__CAG-177
g__Chordicoccus
g__Clostridium
g__Clostridium_AP
g__Collinsella
g__Cryptobacteroides
g__Dorea
g__Eggerthella
g__Gemmiger
g__HGM10890
g__Holdemanella
g__Hominicoprocola
g__Hornefia
g__JAJDHH01
g__Lachnospira
g__Lactobacillus
g__Lentihominibacter
g__Limisoma
g__Limosilactobacillus
g__Megasphaera
g__Oliverpabstia
g__Paenibacillus
g__Paraeggerthella
g__Parafannyhessea
g__Phascolarctobacterium_A
g__Phil12
g__Phocaeicola
g__Prevotella
g__RQZE01
g__RUG023
g__Romboutsia
g__Romboutsia_D
g__Sarcina
g__Sodaliphilus
g__Sphaerochaeta
g__Terrisporobacter
g__Turicibacter
g__UBA1049
g__UBA636
8
6
4
2
0
10301 10302 10303 10701 10702 10703 10801 10802 10803 10901 10902 10903 11001 11002 11003 21001 21002 21003 21101 21102 21103 30101 30102 30103 30601 30602 30603 30701 30702 30703 30801 30802 30803 30901 30902 30903 31101 31102 31103 31201 31202 31203 40501 40502 40503 40601 40602 40603 40901 40902 40903

C.

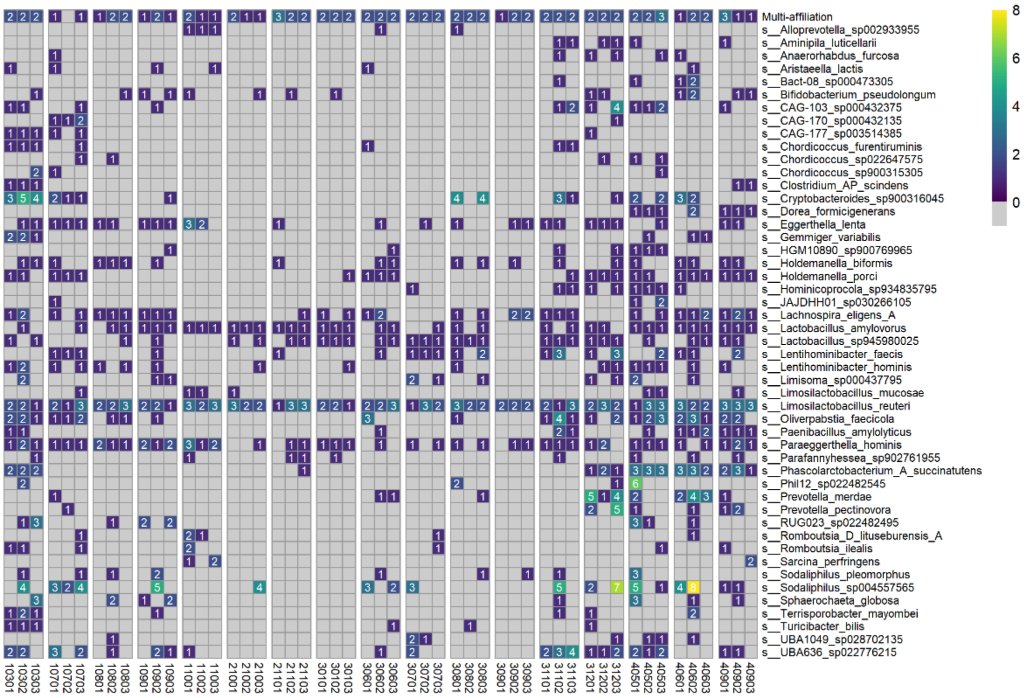

**Figure S7:** Number of G110- or L6-derived sow fecal transmitted and persistent putative strains in L6 and L28 and W5 piglet fecal samples. A. With affiliation at the family level. B. With affiliation at the genus and C. species level. For genus and species affiliation, only the top 50 taxa are presented. (G110: end of gestation, L6: 6 days of lactation, L28: 28 days of lactation, W5: 5 days after weaning))
